# Endogenous Osteoblast-Osteocyte-Osteoclast Signaling Enables Bone Remodeling, Drug Response, and Cancer Invasion in a Nanoscale Calcified Bone-on-a-Chip

**DOI:** 10.64898/2025.12.08.693047

**Authors:** Mauricio G.C. Sousa, Avathamsa Athirasala, Mahshid Hosseini, Daniela M. Roth, Genevieve E. Romanowicz, Rebekka Duhen, Raviraj Thakur, Darya Budkina, May Anny A. Fraga, Sofia M. Vignolo, Aaron Doe, Jinho Lee, Jonathan V. Nguyen, Daquan F. Johnson, Angela S.P. Lin, Eric Orwoll, Cristiane M. Franca, Robert E. Guldberg, Luiz E. Bertassoni

## Abstract

Bone homeostasis depends on spatially orchestrated interactions among osteoclasts, osteoblasts, and osteocytes within a unique microenvironment characterized by an extracellular matrix that is mineralized on the nanoscale. Reconstructing these interactions to enable autonomous cell differentiation and tissue remodeling remains a significant challenge in faithfully modeling bone physiology and disease in-vitro. Here, we present an engineered bone-on-a-chip platform that recapitulates the spatial organization of bone-resident cells, the differentiation of immune cells into osteoclasts, and calcified tissue resorption. We demonstrate that nanoscale mineralization of cell-laden collagen hydrogels is associated with ossification- and matrix-associated transcriptional changes and an osteocyte-like phenotype in osteoblast-lineage cells, while establishing a multicellular microenvironment associated with pro-osteoclastogenic RANKL/OPG signaling and supporting osteoclastogenesis and remodeling without the need for exogenous growth factors. Mineralized niche-driven osteoclastogenesis on-a-chip supported greater osteoclast differentiation than conventional stimulation with RANKL and M-CSF and reproduced differential responses to clinically used anti-resorptive drugs. The model further supports the integration of vascular networks and reproduces features of established tumor-bone interactions, including distinct spatial distributions of different cancer types within the mineralized microenvironment. Together, the findings establish a robust, self-regulated, and versatile microphysiologic model for investigating bone remodeling, cancer-bone crosstalk, and therapeutic interventions.

## 1. Introduction

Bone-related diseases, including fractures, cancers, and osteoporosis, impose a substantial and growing global economic burden^1, 2, 3, 4^. Despite the growing demand for effective bone treatments, existing preclinical models fall short in capturing the complex and dynamic interplay between the cellular machinery and structural components of bone, thus considerably limiting our ability to unravel the mechanisms that govern bone homeostasis, remodeling, and disease^5, 6^. Bone is a dynamic organ system whose formation and homeostasis depend on a tightly orchestrated sequence of cell-cell and cell-matrix interactions occurring within a highly specialized microenvironment^7, 8, 9^. In its mature form, bone is composed of a densely mineralized collagen matrix that embeds cells in three dimensions (3D), forming a stiff, hierarchically organized tissue capable of sensing, adapting to, and remodeling in response to physiological demands.^10, 11, 12^ This mineralization arises through a nanoscale process in which calcium and phosphate ions are precisely guided into collagen fibrils by matrix-regulating proteins, resulting in the formation of intrafibrillar hydroxyapatite crystals^13, 14, 15^. These nanoscale mineralized collagen fibrils define the architecture and mechanical stiffness of the matrix and, critically, establish the cell-binding and ionic domains that regulate cell behavior^16, 17, 18, 19, 20^. For example, it has been demonstrated that cells sense and respond to this mineralized environment through mechanotransductive and ion-mediated signaling pathways that are essential for osteoblast differentiation and osteocyte function^21, 22, 23^. Embedded in the matrix, osteocytes act as central regulators of bone remodeling. They secrete paracrine cues that modulate both osteoblast activity and the recruitment and differentiation of osteoclast precursors^24, 25^. Through this mechanism, osteocytes control the remodeling events that occur in bone tissue, including those involved in resorption, adaptation, and pathologic bone loss^26, 27, 28^. Engineering a human-relevant system that faithfully reproduces these dynamic and endogenously regulated cell-cell and cell-matrix interactions represents not only a complex challenge but a critical need.

Current in-vitro models fail to recapitulate many of the essential hallmarks of bone, including chemical, physical, and biological features^29^. Most rely on non-mineralized collagen matrices or apatite-laden hydrogels that lack the biochemical complexity and mechanical cues of the native nanoscale-calcified bone matrix. Alternatives such as mineral-coated plates, bone explants, and demineralized bone surfaces, though considered gold standards for studying bone resorption, replicate some of the properties of native bone while neglecting the complex interplay of cell-cell and cell-matrix communication that is known to result in a self-regulating remodeling unit that defines bone physiology^30, 31, 32^. For example, osteoclastogenesis in these models is typically induced through supraphysiologic supplementation with recombinant receptor activator of nuclear factor κB ligand (RANKL) and macrophage colony-stimulating factor (M-CSF), thereby bypassing the natural signaling hierarchy regulated by the communication between osteocytes, osteoblasts, and osteoclasts^33, 34^. Even advanced 3D and organ-on-a-chip platforms often lack the combination of true bone-like nanoscale mineralized ECM, spatially organized bone cell populations, and endogenous signaling necessary to reproduce coordinated bone remodeling.^35, 36^. These limitations restrict our ability to investigate how matrix mineralization and local cellular interactions jointly regulate osteoclastogenesis, tissue resorption, and responses to disease or therapeutic interventions.

To address these limitations, here we introduce a bone-on-a-chip platform that combines nanoscale intrafibrillarly mineralized collagen with spatially organized osteoblast-lineage cells, osteoclast precursors, and endothelial cells. Within this engineered microenvironment, matrix mineralization promotes a mineral-responsive, osteocyte-like state of osteoblast-lineage cells and functional osteoclastogenesis through endogenous cell-cell and cell-matrix communication without the need for exogenous supplements and growth factors. Moreover, this self-regulated system allows for osteoclast differentiation events with distinct associated transcriptional profiles from those observed with standard M-CSF- and RANKL-supplemented differentiation medium. Mineralized niche-driven osteoclastogenesis exhibits distinct secretory and transcriptional profiles, with enrichment of matrix-associated adhesion, migration, and remodeling programs, compared with the inflammatory signaling programs preferentially enriched following conventional RANKL and M-CSF stimulation. By maintaining the mineralized matrix, bone-resident cells, and the osteoclast precursors in spatially defined compartments, the platform enables investigation of how local cell-cell and cell-matrix interactions influence the RANKL/OPG axis, osteoclast differentiation, and activity. The platform also accommodates vascular network formation, incorporating an essential structural component of native bone and extending its applicability to the study of vascular and cancer–bone interactions. The resulting system exhibits robust resorptive behavior observed via micro-CT, responds to clinical-grade anti-resorptive drugs, and models cancer-bone interactions, including direct bone invasion by oral squamous cell carcinoma cells and the vascular association of prostate cancer cells. In summary, this biomimetic bone-on-a-chip platform addresses critical gaps in existing models by authentically replicating the complex microenvironment of bone. This provides a novel and robust, physiologically relevant new approach methodology (NAM) tool to study bone homeostasis, disease mechanisms, and therapeutic responses in-vitro for preclinical research in bone-targeting diseases and therapeutic interventions.

## 2. Results

### 2.1. Nanoscale mineralization of an engineered multicellular bone microenvironment promotes an osteocyte-like phenotype

Bone is a hierarchical structure organized at the nanoscale level by intrafibrillarly mineralized collagen fibers, which are critical for osteocyte differentiation and the regulation of osteoclastogenic factors^37^. To replicate this nanostructure in-vitro, we developed a biomimetic model by embedding osteoblasts in a collagen matrix (2.5 mg/mL) within a microfluidic device featuring a central channel flanked by two lateral channels that can be perfused independently (**Figure 1a**). The osteoblast-laden collagen matrix was injected into the main channel and underwent mineralization over a 3-day period. We followed a previously developed strategy to achieve mineralization using previously optimized media containing physiological levels of calcium (Ca^2+^), phosphate (PO ^3-^), and a matrix protein analogue (Lacprodan, mOPN-10). This media composition delays premature precipitation of the calcium and phosphate in solution, allowing amorphous calcium phosphate to penetrate individual collagen fibrils and nucleate intrafibrillarly, closely mimicking native mechanisms of bone mineralization as demonstrated in previous studies^38, 39^. As a result, the engineered matrix recapitulates both the composition and hierarchical organization of the native bone tissue.

**Figure 1.**
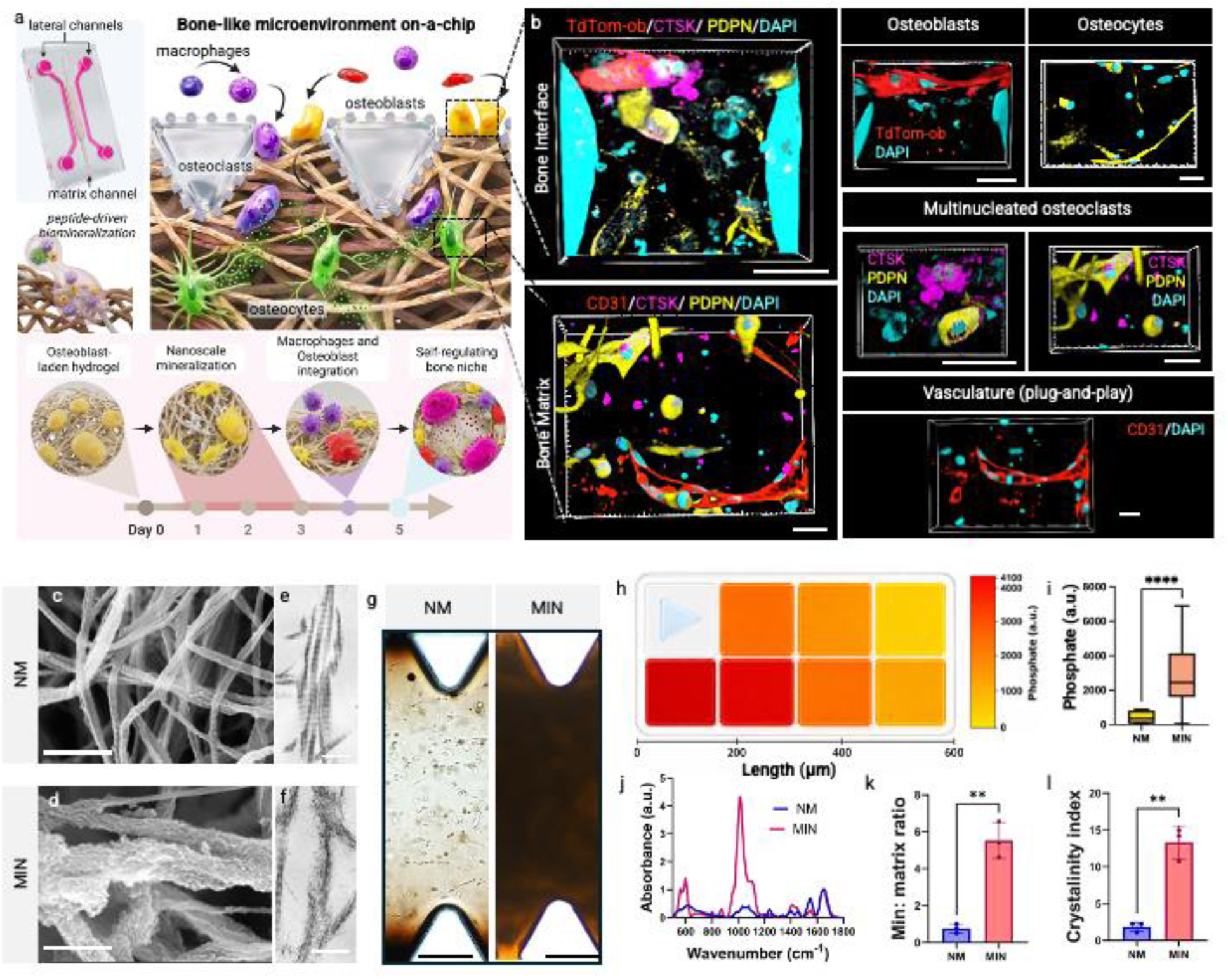
A bone-on-a-chip that recapitulates the self-regulating multicellular complexity of bone within a mineralized matrix. (a) Conceptual schematic of the organ-on-a-chip platform, illustrating the stepwise engineering of a mineralized bone-like microenvironment that establishes the physiological spatial organization of bone cells and recapitulates key in-vivo interactions. (b) The engineered system supports the co-existence and direct interaction of osteoblasts (tdTomato; red), osteocyte-like cells (PDPN; yellow), and osteoclasts (CTSK; magenta), established without exogenous differentiation factors. In a separate configuration, the platform integrates a vascular network, shown with osteocyte-like cells (PDPN) and endothelial vessels (CD31; red). Scale bars 30 µm. (c–d) Scanning electron microscopy and (e–f) transmission electron microscopy of non-mineralized (NM; c, e) and mineralized (MIN; d, f) matrices reveal discrete mineral deposits and increased apparent fibril thickness in mineralized samples, consistent with mineral deposition associated with the fibrillar network. Scale bars 250 nm. (g) Alizarin Red staining of non-mineralized and mineralized conditions confirms calcium deposition in the mineralized matrix. Scale bars 200 µm (h) Confocal Raman mapping of phosphate distribution (ν₁PO₄, a.u.) across the mineralized construct, from the channel edge toward the center, showing higher phosphate signal near the pillars and a gradient toward the core on a color scale. (i) Quantification of the phosphate Raman signal (a.u.) in non-mineralized versus mineralized samples with bar colors representing the same color scale showed significantly greater phosphate content in mineralized constructs. (j) Fourier-transform infrared (FTIR) spectra, (k) mineral-to-matrix ratio, and (l) crystallinity index further confirm increased mineral content in the mineralized group. All experiments were performed with three biological replicates. Statistical significance in (i), (k), and (l) was assessed by two-tailed Student’s t-test (****p < 0.0001, **p < 0.01). Scale bars in b represent 30 µm. Parts of this figure were created with BioRender.

Beyond recreating the mineralized matrix, the platform enables the sequential incorporation and spatial organization of multiple bone-associated cell populations (**Figure 1a-b**). Osteoblasts embedded within the mineralized matrix exhibit osteocyte-associated characteristics and become entrapped within the calcified collagen. Subsequent addition of osteoblasts and macrophages in the lateral channels resulted in their lining the tissue interface, where the macrophages differentiated into osteoclasts. Together, these features establish a self-organizing, physiologically relevant bone-like microenvironment governed by endogenous cell–cell and cell–matrix interactions encoded by the platform design. We further demonstrate the feasibility of adapting the platform to model complex, vasculature-driven multicellular interactions by incorporating an endothelial network that interfaces with the mineralized bone-like matrix, thereby resulting in a multicellular system consisting of vasculature, osteocytes, and osteoclasts (**Figure 1b, Supporting video 1**). Notably, quantitative analysis revealed no significant differences in total vessel length or area between mineralized and non-mineralized conditions, indicating that mineralization did not impair vascular morphogenesis (**Supplementary Figure S1**). Importantly, this modular approach enables the stepwise integration of relevant tissue components, such as vasculature, to facilitate investigations of how vascular cues contribute to bone physiology and disease progression.

To validate the formation of a bone-like matrix, we assessed nanoscale mineralization relative to non-mineralized control samples (NM). Clear differences in opacity were observed between the non-mineralized and mineralized samples (MIN) by light microscopy (**Supplementary Figure S2a-b**), which were attributable to mineralization, as verified by Alizarin red staining, which showed visible evidence of mineral deposition in the mineralized groups (**Figure 1g**). Scanning electron microscopy (SEM) analysis (**Figure 1c-d**) revealed pronounced differences in fiber structure between the groups, with distinct mineral deposits on the collagen fibers apparent in the mineralized samples. Moreover, the collagen fibers in the mineralized matrices appeared visibly thicker and were accompanied by surface roughness (**Figure 1d**). These structural changes, however, were notably absent in the non-mineralized samples (**Figure 1c**), offering further evidence supporting successful incorporation of nanoscale mineral in the collagen matrix. Transmission electron microscopy (TEM) further resolved these differences at the fibrillar level, with mineralized samples exhibiting electron-dense apatite crystallites associated with collagen fibrils, consistent with collagen intrafibrillar mineralization (**Figure 1f**), which were absent in non-mineralized controls (**Figure 1e**). Importantly, cell viability remained high following mineralization, with comparable viability observed in MIN and NM constructs (**Supplementary Figure S2c–e**). Confocal Raman spectroscopy was used to map the spatial distribution of mineral across the construct. Phosphate signal (ν₁ PO₄^3-^) was detectable throughout the mineralized matrix, with higher intensity near the channel edges gradually decreasing toward the center (**Figure 1h**). Even in the central region with the lowest signal, phosphate intensity remained well above that of non-mineralized controls, confirming that mineralization occurred throughout the construct despite the spatial gradient. Spatially resolved Raman spectra revealed phosphate-associated signals across the sampled matrix regions, with greater intensity in MIN than in NM constructs (**Supplementary Figure S3**). Quantification confirmed significantly greater phosphate signal in mineralized samples compared to non-mineralized controls across all positions sampled (**Figure 1i**). Additionally, Fourier Transform Infrared Spectroscopy (FTIR) was performed to confirm the chemical composition of the mineralized matrix. The mineralized samples exhibited prominent peaks corresponding to expected apatite phosphate (1030, 600, and 560 cm⁻¹) and carbonate (874 cm⁻¹), absent in the non-mineralized groups (**Figure 1j**). These findings align with patterns observed in native bone^38, 39, 40^. Quantitative analyses showed significantly higher mineral-to-matrix ratios (**Figure 1k**) and crystallinity indices (**Figure 1l**) in mineralized samples compared to controls, and comparable mineralization relative to native bone^40^, corroborating the ability of this model to replicate key properties of bone tissue at the nanoscale level.

To determine how the mineralized matrix influences osteoblast differentiation, we first performed bulk RNA sequencing of mineralized and non-mineralized osteoblast-laden collagen matrices cultured off-chip and harvested at day 3 (n = 4). Principal component analysis revealed clear separation between MIN and NM groups primarily along PC1, indicating distinct transcriptional profiles associated with mineralization (**Supplementary Figure S4a**). Differential expression analysis identified 2864 significantly differentially expressed genes, including 1401 upregulated and 1463 downregulated in MIN relative to NM (adjusted p < 0.05). Among the most strongly upregulated genes in MIN were *CRLF1*, *SERPINE1*, *LBH*, and *CDH2*, which have been implicated in skeletal development, extracellular matrix remodeling, and cell adhesion^41, 42, 43, 44, 45^, whereas *STAT1* and *MX2*, both associated with interferon signaling, were higher in NM (**Figure 2a**). Analysis of the top differentially expressed genes further highlighted transcriptional changes associated with ECM remodeling, cytoskeletal organization, and developmental signaling in MIN, in contrast to interferon-responsive and sterol-biosynthetic signatures in NM (**Supplementary Figure S4b,c**). Gene set enrichment analysis showed that ossification, skeletal system morphogenesis, cytoskeletal organization, bone development, bone mineralization, and osteoblast and osteoclast differentiation were enriched in MIN, whereas negative regulation of osteoblast proliferation was enriched in NM (**Figure 2c**).

**Figure 2.**
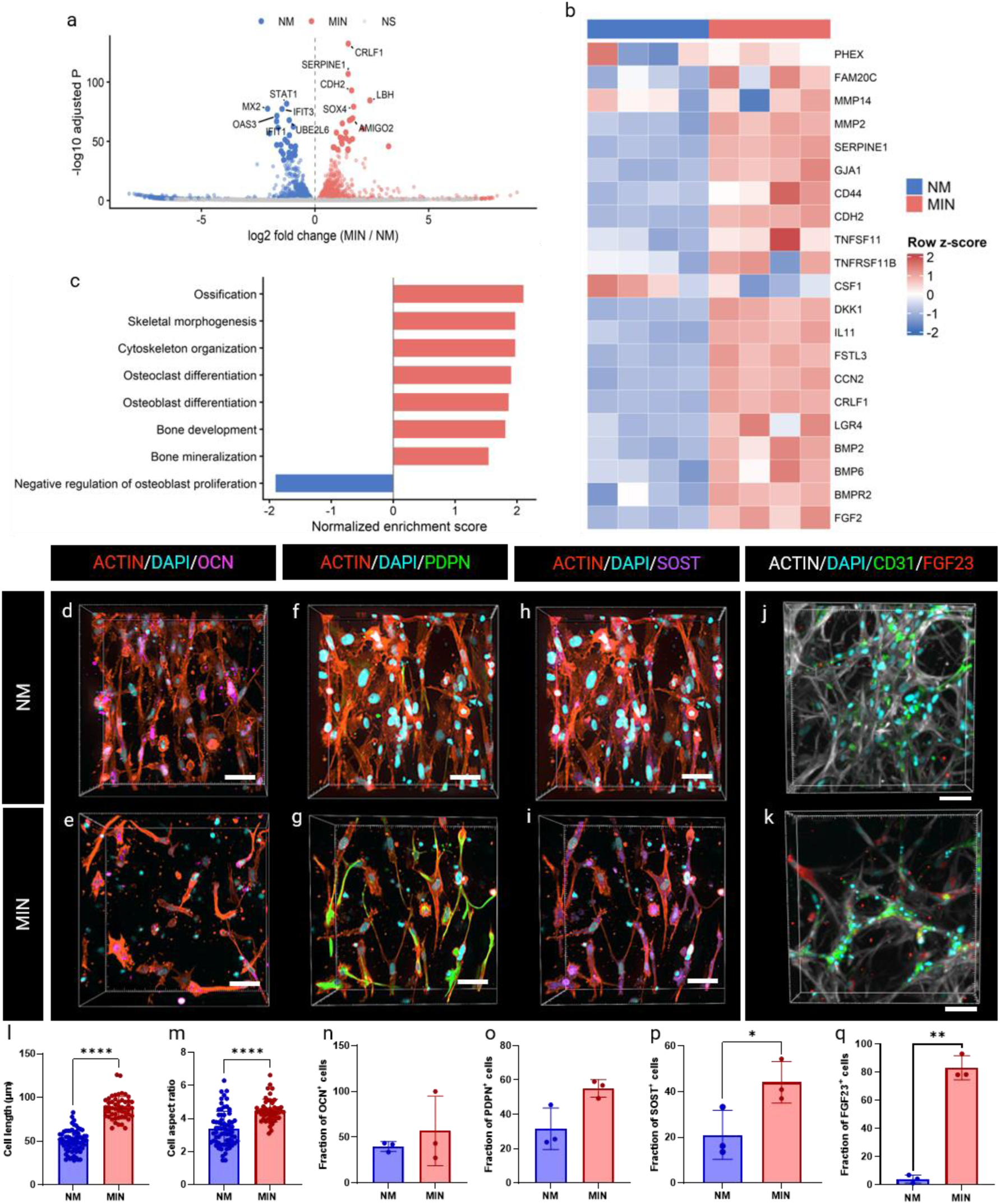
Matrix mineralization is associated with distinct osteoblast-lineage transcriptional profiles and osteocyte-like phenotypic changes. (a) Volcano plot of bulk RNA-seq comparing osteoblast-laden MIN and NM collagen constructs cultured off-chip for 3 days (n = 4) showing differentially expressed genes (DESeq2, p < 0.05), Genes higher in MIN include CRLF1, LBH, SERPINE1, and CDH2, whereas STAT1 and MX2 are higher in NM. (b) Heatmap of selected genes associated with bone matrix maturation, mineralization, and osteoblast-lineage signaling, displayed as row-normalized z-scores, with red indicating higher and blue indicating lower relative expression. (c) Preranked gene set enrichment analysis (GSEA) performed using fgsea, with genes ranked by the DESeq2 Wald statistic, showing normalized enrichment scores (NES) for selected biological processes (FDR < 0.05) with positive and negative NES scores indicating enrichment in MIN and NM respectively. (d–i) Representative confocal images of osteoblast-lineage cells in NM and MIN chips stained for F-actin (red), DAPI (blue), and OCN (magenta), PDPN (green), or SOST (magenta), illustrating changes in (l,m) cellular morphology of individual cells and (n,o,p) osteocyte-associated marker expression following matrix mineralization. (j, k) Representative confocal images of vascularized constructs incorporating primary human osteoblasts, stained for F-actin (white), DAPI (blue), CD31 (green), and FGF23 (red) showed (q) higher percentage of FGF23 positive cells in mineralized conditions in the vascularized primary-osteoblast constructs. Data represent mean ± SD (standard deviation). (n = 3 biological replicates for panels l–p and q). Statistical significance was determined using two-tailed unpaired Student’s t-tests, except for FGF23 positivity, which was analyzed using a linear mixed-effects model with Kenward–Roger-adjusted pairwise comparisons and Holm correction for multiple testing (*p < 0.05, **p < 0.01, ***p < 0.001, ****p < 0.0001). Scale bars: 50 µm (d–k).

To further investigate the influence of matrix mineralization on bone-associated signaling, we examined a curated set of genes involved in matrix mineralization, remodeling, cell adhesion, and intercellular communication (**Figure 2b**). MIN samples exhibited higher expression of genes including *PHEX, MMP14, SERPINE1, CDH2, DKK1, BMP2, BMP6*, and *BMPR2*, suggesting changes in matrix regulation and intercellular signaling. *TNFSF11* (RANKL) and *TNFRSF11B* (OPG)^46^ also exhibited higher expression in MIN, whereas *CSF1* (M-CSF) was lower, indicating that mineralization is associated with changes in the transcriptional profile of key regulators of osteoclastogenesis. These findings suggest that the mineralized matrix may influence the signaling environment associated with osteoclast regulation. Further examination of transcriptional regulators revealed increased expression of *ATF4*^47^ and several AP-1 family members, including *FOS, FOSB, JUN, JUNB*, and *JUND* in MIN, consistent with a transcriptional response associated with the mineralized matrix (**Supplementary Figure S4d**). In contrast, genes associated with osteoblast differentiation exhibited a less coordinated response, with variable expression of lineage-associated markers such as *RUNX2, DLX5*, and *ALPL*, while genes involved in matrix organization and cellular interactions, including *CCN1, VCAN, TNC*, and *CDH11*, were elevated in MIN (**Supplementary Figure S4e**). Together, these findings suggest that non-mineralized conditions retain aspects of an earlier osteoblast differentiation program, whereas matrix mineralization promotes a distinct transcriptional response consistent with progression toward an osteocyte-like phenotype.^48^

In the physiological context of bone, osteoblasts secrete a dense collagenous matrix, termed osteoid, which eventually mineralizes to entrap the cells, triggering their transition into osteocytes^49^. To determine whether this transcriptional response was accompanied by an osteocyte-like cellular phenotype, we examined the morphology and protein expression of osteoblast-lineage cells in the chips on day 3. In the mineralized groups, cells displayed marked morphological remodeling consistent with an osteocyte-like phenotype, including elongated cell bodies, increased aspect ratios, and the development of dendritic cytoplasmic extensions forming an interconnected network (**Figure 2l-m and Supplementary Figure S5b**). In contrast, cells in non-mineralized conditions retained a rounded morphology with limited projections, consistent with an osteoblastic state (**Supplementary Figure S5a**). Mineralization also altered the abundance and distribution of proteins associated with the osteocyte phenotype and function. Immunostaining revealed a trend toward increased osteocalcin (OCN) and podoplanin (PDPN) positivity in MIN that did not reach statistical significance, although PDPN localization along cellular extensions was consistent with the development of an osteocyte-like morphology (**Figure 2d-g and n-o**). The fraction of cells positive for sclerostin (SOST), a marker associated with late osteocyte differentiation, was significantly higher in the mineralized group (**Figure 2h-i**, **p**). In contrast, the fraction of cells positive for DMP1 and FGF23 did not differ significantly between mineralized and non-mineralized conditions (**Supplementary Figure S5c-f and k-l**).

Together, these findings indicate that the mineralized microenvironment promotes coordinated structural changes and protein expression patterns consistent with osteocyte-lineage progression ^37, 50^. Collectively, the elongation of the osteoblasts, the formation of dendritic extensions, the higher number of SOST positive cells, and the PDPN localization along dendrites are compatible with a partial shift from an osteoblastic toward an osteocyte-like phenotype^51, 52^. To further evaluate the osteocyte-associated phenotype induced by matrix mineralization, we compared protein expression with cultures treated with conventional osteoinductive medium (OIM) for three days. The fractions of OCN- and PDPN-positive cells did not differ significantly between MIN and OIM conditions, whereas the fraction of SOST-positive cells was significantly higher following matrix mineralization (**Supplementary Figure S5g-i and m-o**). These data indicate that matrix mineralization promotes osteocyte-associated marker positivity comparable to or greater than that observed with biochemical osteoinduction under comparable culture conditions. To further assess the applicability of the platform, we incorporated primary human osteoblasts into vascularized bone-on-a-chip constructs. In this configuration, mineralization significantly increased the fraction of FGF23-positive primary human osteoblast cells compared with non-mineralized controls (**Figure 2j–k,q),** demonstrating the capacity of the platform to support osteocyte-associated protein expression in a more complex multicellular environment.

These phenotypic changes occurred under conditions of low predicted interstitial fluid shear. Our computational fluid dynamics simulation estimated an interstitial velocity of ∼1 µm/s and a shear stress of ∼0.001 Pa in the non-mineralized collagen, well below the 0.5–5 Pa reported to drive osteoblast differentiation in-vitro (**Supplementary Figure S6**). Because even this presumed more permeable condition fell far below that range, the transition is unlikely to be primarily driven by fluid shear, supporting a role for matrix mineralization in promoting the observed phenotypic changes. Furthermore, both MIN and NM constructs were maintained under identical rocking conditions, indicating that differences in the observed cellular phenotypes cannot be attributed to differences in the externally imposed flow conditions. Together, these findings support a predominant role for matrix mineralization in promoting the observed osteocyte-associated phenotypic changes^53^. This mineral-responsive state emerges within days and, together with the transcriptional program described above, supports the development of a bone-prototypical environment for remodeling.

### 2.2. Mineralized matrix and multicellular crosstalk support functional osteoclastogenesis

Osteocytes are central regulators of bone homeostasis, coordinating osteoblast and osteoclast activity through key signaling pathways, including the modulation of the ratio between receptor activator of nuclear factor κB ligand (RANKL) and osteoprotegerin (OPG)^54^, in which RANKL promotes osteoclastogenesis, and OPG serves as its inhibitor (**Figure 3a**). These cues, influenced by mechanical factors like matrix stiffness and fluid shear stress, bridge biomechanical and biochemical cues, facilitating the recruitment of monocytes and their differentiation into osteoclasts^55, 56^. While these interactions are well-characterized in-vivo, replicating them in-vitro remains challenging, particularly when sustaining multiple differentiated cell types in culture using exogenous growth factors alone, which can be inefficient and lead to unintended crosstalk. In our system, following matrix mineralization, which induced the mineral-responsive program described above, we leveraged the resulting microenvironment to spatially recreate the coordinated triad of bone cell activity. To do so, we introduced additional osteoblasts followed by human monocyte-derived macrophages, either isolated from patient peripheral blood or from a human cell line, into the lateral channel of the microfluidic chip. The added osteoblasts were intended to line the mineralized matrix surface, while the osteocyte-like cells remained embedded within the mineralized matrix. The macrophages served as osteoclast precursors at the interface, thereby reconstituting the three principal bone cell types within a single mineralized niche. This configuration simulates the physiological recruitment of monocytes/macrophages to the bone surface, where they undergo osteoclastogenesis on a mineralized microenvironment (**Figure 3a**). The osteoclast phenotype in mineralized chips was compared with control samples either left untreated or treated with RANKL (50 ng/mL) and macrophage colony-stimulating factor (M-CSF, 30 ng/mL), representing the conventional growth factor-mediated osteoclast differentiation protocol (named NM + GF)^57^.

**Figure 3.**
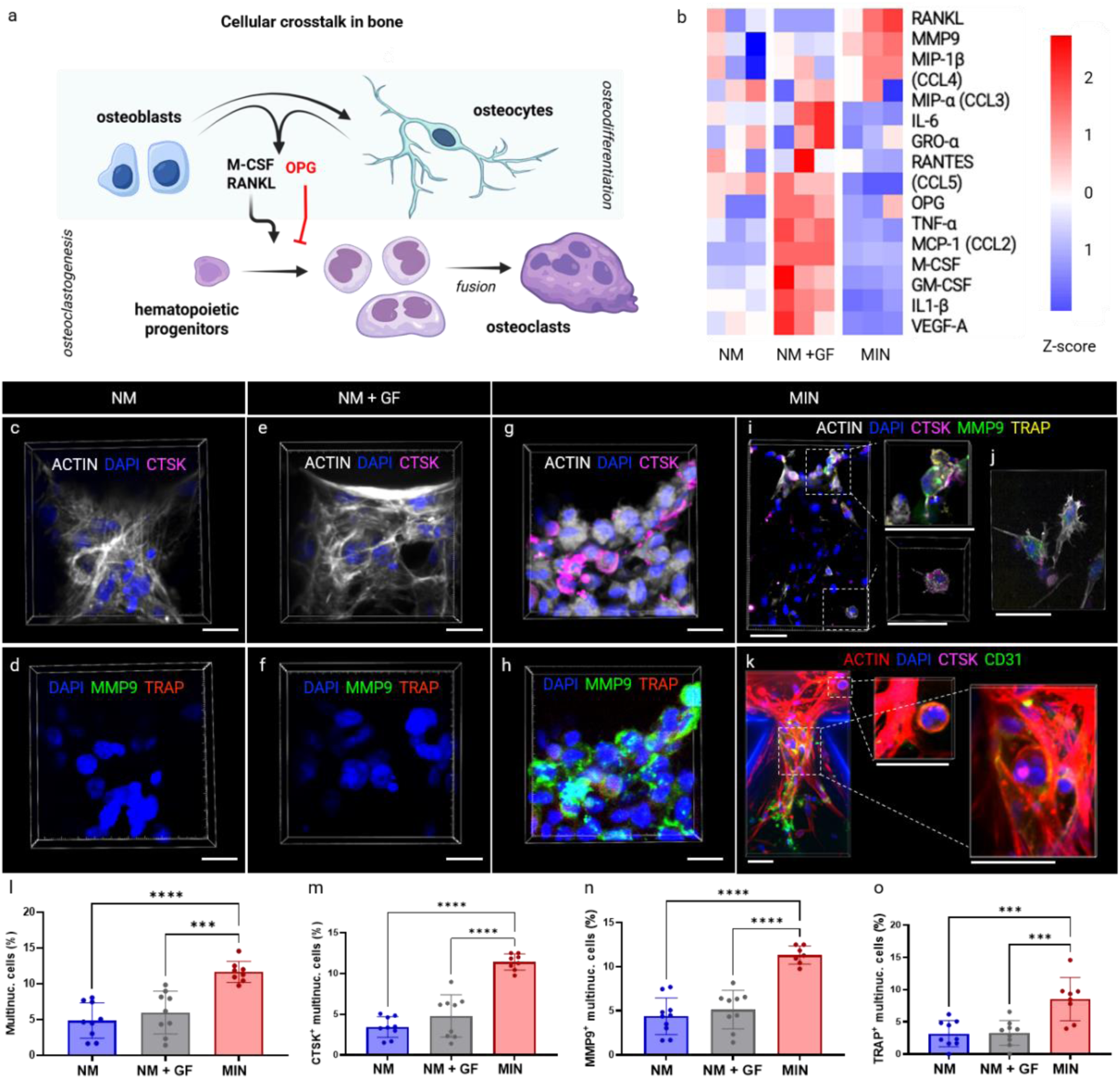
The mineralized bone-on-a-chip supports osteoclastogenesis without exogenous RANKL or M-CSF supplementation. (a) Schematic illustration of cellular crosstalk between osteoblast-lineage cells and osteoclast precursors, highlighting the roles of RANKL, M-CSF, and (OPG) in regulating osteoclastogenesis. (b) Heatmap of cytokines and growth factors measured by multiplex Luminex analysis of day 3 culture supernatants from non-mineralized constructs (NM), non-mineralized constructs supplemented with RANKL (50 ng mL⁻¹) and M-CSF (30 ng mL⁻¹) (NM + GF), and mineralized constructs without exogenous RANKL or M-CSF (MIN). Exogenously supplemented RANKL and M-CSF concentrations were subtracted from the corresponding NM + GF measurements before analysis. Row-normalized z-scores illustrate distinct secretory profiles, with higher RANKL and lower OPG in MIN compared with NM + GF, whereas growth factor supplementation was associated with increased inflammatory cytokine secretion. (c–h) Representative day 5 confocal images of macrophage-derived osteoclast-associated cells in NM (c, d), NM + GF (e, f), and MIN (g, h), stained for F-actin (white), DAPI (blue), and cathepsin K (CTSK; magenta) (c, e, g), or DAPI (blue), (MMP9; green), and (TRAP; red) (d, f, h), showing increased multinucleation and osteoclast-associated marker expression in MIN. (i) Representative confocal images showing qualitative differences in osteoclast-associated cell morphology and marker expression according to spatial location within the lateral channel and mineralized matrix. (j) Representative images of primary human monocyte-derived osteoclast-associated cells in mineralized constructs, demonstrating multinucleation and osteoclast-associated marker expression without exogenous RANKL or M-CSF during on-chip culture. (k) Representative confocal images of vascularized mineralized constructs incorporating primary human osteoblasts, THP-1-derived macrophages, and endothelial cells, showing CTSK-positive cells in proximity to CD31-positive vascular structures. (l–o) Quantification of multinucleated cells (l), CTSK-positive multinucleated cells (m), MMP9-positive multinucleated cells (n), and TRAP-positive multinucleated cells (o), demonstrating increased osteoclastogenesis in MIN compared with non-mineralized conditions. Multinucleated cells were defined as cells containing three or more nuclei. Data represent mean ± SD from three biological replicates per condition. Statistical significance was determined by ordinary one-way ANOVA followed by Tukey’s multiple-comparisons test (***p < 0.001, ****p < 0.0001). Scale bars: 20 µm (c–h) and 50 µm (i–k).

To investigate the signaling mechanisms underlying osteoclastogenesis within the engineered bone microenvironment, we performed multiplex Luminex analysis of secreted cytokines and growth factors across the experimental conditions. Distinct secretory profiles emerged between mineralized constructs and non-mineralized controls, with or without conventional osteoclastogenic supplementation (**Figure 3b**). By day 3, mineralized constructs exhibited increased RANKL levels, accompanied by lower OPG levels relative to growth factor-supplemented controls, consistent with a signaling environment favorable to osteoclastogenesis (**Supplementary Figure S8i,k**). Differences were also observed in osteoclast-associated mediators, including MMP9 and macrophage inflammatory protein-1α (MIP-1α/CCL3). For this comparison, the concentrations of RANKL and M-CSF that were added to the NM + GF medium were subtracted before comparing groups; after this correction, the mineralized samples showed higher RANKL and lower OPG and M-CSF levels. In contrast, conventional RANKL and M-CSF supplementation was associated with elevated levels of inflammatory mediators, including IL-6, TNF-α, and MCP-1 (CCL2). These findings indicate that the mineralized niche establishes a distinct secretory environment that supports osteoclastogenesis without exogenous osteoclastogenic stimulation.

Longitudinal analysis further revealed distinct temporal patterns of cytokine secretion across the experimental conditions (**Supplementary Figures S7 and S8**). In mineralized groups, RANKL levels peaked at day 3 but returned to baseline by day 5, while OPG remained comparatively throughout the experimental period. On the other hand, conventional RANKL and M-CSF stimulation elicited a broader inflammatory secretory response, characterized by elevated TNF-α and GM-CSF at early time points and comparatively higher IL-6, IL-1β, and MCP-1 levels at later time points. Growth factor-supplemented constructs also exhibited higher VEGF secretion, particularly at day 5. Together, these findings suggest that the mineralized niche supports a temporally distinct osteoclast-associated signaling environment compared with conventional growth factor supplementation. Together, these findings suggest that the mineralized niche supports a temporally distinct osteoclast-associated signaling environment compared with conventional growth factor supplementation. Interestingly, *TNFRSF11B* (OPG) transcript levels were higher in mineralized osteoblast-lineage cultures in the day-3 RNA-seq analysis (**Figure 2b**), whereas secreted OPG protein levels were lower in the multicellular mineralized constructs (**Figure 3b**). These differences may reflect the distinct cellular compositions and experimental configurations, as well as differences between transcript abundance and protein secretion. We therefore base our interpretation of the pro-osteoclastogenic RANKL/OPG balance on the secreted protein measurements. Because the Luminex measurements were obtained from bulk supernatant of chips in which osteoblast-lineage cells are present in more than one channel, we do not assign the cellular source of RANKL and instead attribute this response to the multicellular crosstalk between osteoblasts, osteocytes and osteoclasts in the presence of the mineralized niche.

Next, we evaluated osteoclast differentiation within the engineered bone microenvironment. Ranging from days 3 to 5, multinucleated osteoclasts derived from a human monocytic cell line expressed cathepsin K, MMP9, and TRAP which were significantly more abundant in mineralized conditions compared to non-mineralized controls, irrespective of exogenous RANKL and M-CSF supplementation (**Figure 3c–h, l-o**). Supplementation of mineralized samples with RANKL and M-CSF (MIN + GF) further increased the proportion of cathepsin K-, TRAP-, and MMP9-positive multinucleated cells (**Supplementary Figure S9**), demonstrating the tunability of the system for modeling graded and potentially individualized bone remodeling responses. Notably, in conventional 2D culture systems, osteoclastogenesis typically requires approximately 14 days or more of continuous RANKL/M-CSF stimulation. In our platform, incorporation of the biomechanical and biochemical cues secreted by bone cells embedded in a mineralized matrix supported osteoclast differentiation over a substantially shorter period. To investigate whether osteoclastogenesis could be reproduced by individual components of the engineered microenvironment, we performed additional control experiments. Neither bovine milk-derived osteopontin supplementation of non-mineralized, cell-laden matrices nor an acellular mineralized matrix reproduced the osteoclastogenic response observed in the complete mineralized constructs, as assessed by the formation of multinucleated cells expressing CTSK and MMP9 (**Supplementary Figure S11**). Similarly, conditioned medium collected from mineralized constructs did not induce appreciable osteoclast differentiation in macrophages cultured in two dimensions (2D), whereas conventional RANKL/M-CSF stimulation for 14 days confirmed their osteoclastogenic potential (**Supplementary Figure S10**). Together, these findings suggest that robust osteoclastogenesis depends on the combined cellular and matrix-associated cues of the mineralized microenvironment, which were not reproduced by the individual conditions tested in isolation.

In addition to enhanced osteoclast differentiation, we observed spatial differences in the morphology and distribution of osteoclast-associated markers within the construct. Cells within the lateral channels exhibited predominantly rounded, multinucleated morphologies, whereas those extending into the mineralized matrix appeared more elongated, with cytoplasmic projections (**Figure 3i**). Qualitative immunostaining also suggested more prominent TRAP and MMP9 staining in cells within the lateral channels compared with those located within the matrix. These observations suggest potential spatial heterogeneity in osteoclast-associated phenotypes within the engineered bone microenvironment.

To confirm that this response was not specific to the monocytic cell line, we repeated the experiments using monocytes isolated from peripheral blood mononuclear cells of healthy donors (**Supplementary Figure S12a-c**). Following differentiation into macrophages ex-vivo, cells were introduced into the lateral channels of the chips (**Supplementary Figure S12a**). Consistent with the cell line findings, mineralized constructs supported significantly greater proportions of multinucleated cells expressing CTSK and MMP9 compared with non-mineralized controls, even in the absence of additional RANKL and M-CSF supplementation (**Supplementary Figure S12d,e**). Growth factor supplementation further enhanced this response. These findings demonstrate that the osteoclastogenic effects of the mineralized microenvironment extend to primary human monocyte-derived macrophages.

We further evaluated the versatility of the platform by incorporating primary human osteoblasts, endothelial networks, and THP-1-derived macrophages into a vascularized bone-like microenvironment. CTSK-positive osteoclast-associated cells were observed in close proximity to CD31-positive vascular structures and the mineralized tissue interface (**Figure 3k, Supporting Video 1**). These observations demonstrate the compatibility of osteoclast-like cells with vascular networks and primary osteoblasts within the engineered microenvironment, providing a foundation for investigating more complex interactions among bone-resident, vascular, and immune cells.

### 2.3. Distinct transcriptional programs accompany mineralized niche-driven and conventional osteoclastogenesis

To elucidate the molecular programs associated with the biomimetic osteoclastogenesis observed in **Figure 3**, we performed targeted gene expression profiling to compare differentiating cells exposed to mineralized versus growth factor-supplemented non-mineralized matrices. Osteoclastogenesis is classically regulated by the RANKL/RANK and M-CSF signaling pathways, which converge on activation of NF-κB to initiate transcriptional programs associated with osteoclast differentiation and activity^58^. While these pathways are well-characterized in 2D cultures supplemented with exogenous RANKL and M-CSF, the signaling mechanisms engaged by a mineralized cell-laden matrix remain poorly explored in-vitro^57^. Our platform provides a unique opportunity to dissect these pathways that arise from multicellular crosstalk within the mineralized microenvironment.

To evaluate how our engineered mineralized matrix compares to the traditional growth factor-driven method of osteoclastogenesis, we compared the gene expression profiles of cells recovered mainly from the lateral channel (osteoclast precursors and osteoblasts, with a minor contribution of matrix-resident cells) of biomimetically mineralized chips with those on standard collagen matrices supplemented with exogenous RANKL/M-CSF (**Figure 4**). The non-mineralized, growth factor-free condition (NM) was also included as a baseline, allowing the transcriptional profiles of both osteoclastogenic conditions to be evaluated relative to unstimulated controls (**Supplementary Figure S13c**). RNA isolated after three days of incubation following introduction of macrophages was analyzed using a NanoString nCounter panel targeting 825 genes (**Supplementary Table 1**). Principal component analysis revealed clear separation between MIN and NM + GF conditions along PC1, accounting for 65% of the variance (**Supplementary Figure S13a**). Both groups exhibited enrichment of classical osteoclast markers, such as *ACP5 (TRAP)* and *CTSK* (**Supplementary Figure S13b**). Differential expression analysis identified distinct transcriptional profiles, with inflammatory-associated genes, including IL1B, STAT1, and FSTL1, elevated in NM + GF, whereas genes including CST7, FBLN5, CLDN3, and ADAM8 were more highly expressed in MIN (**Figure 4a**).

**Figure 4.**
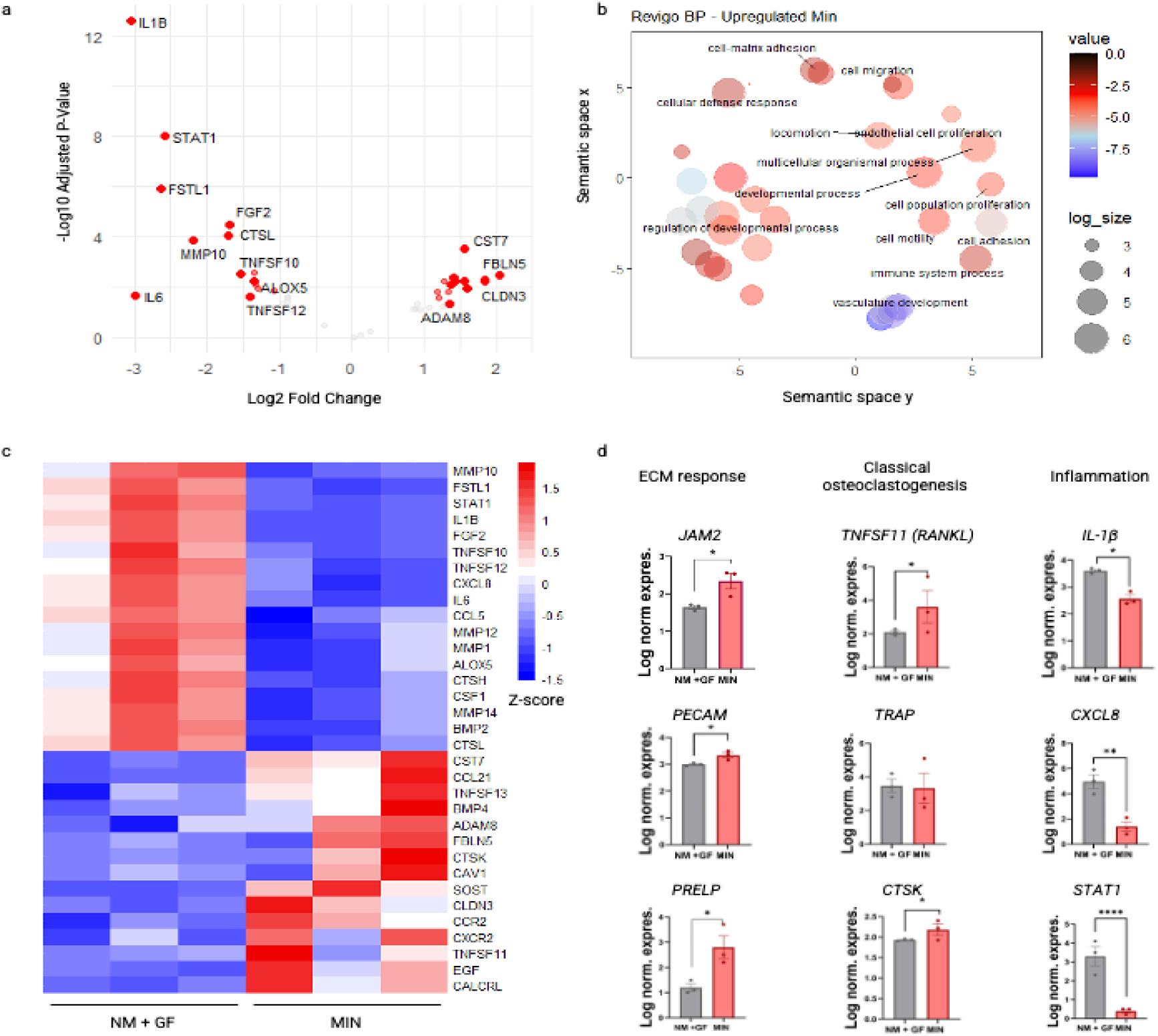
Distinct transcriptional signatures accompany osteoclastogenesis within the mineralized bone microenvironment. Targeted NanoString analysis of cells collected predominantly from the lateral channels of mineralized constructs (MIN) and non-mineralized constructs supplemented with RANKL and M-CSF (NM + GF) at day 3 revealed distinct transcriptional profiles associated with cell–matrix interactions, migration, and inflammatory signaling. (a) Volcano plot showing differential gene expression between MIN and NM + GF using an 825-gene NanoString panel. Each point represents a gene plotted according to its log₂ fold change (MIN/NM + GF) and adjusted p-value (p˂0.01), with red indicating significantly differentially expressed genes. (b) ReviGO visualization of Gene Ontology (GO) biological processes enriched among the top 100 genes upregulated in MIN relative to NM + GF, highlighting processes associated with cell–matrix adhesion, migration, motility, and cellular organization. GO enrichment analysis was performed using the Princeton GO Term Finder, with biological process terms meeting p < 0.1 included in the visualization. (c) Heatmap showing row-normalized z-scores of a curated set of genes associated with extracellular matrix interactions, osteoclast-related signaling, and inflammation in MIN and NM + GF constructs. Red indicates higher and blue indicates lower relative expression, with each column representing an individual biological replicate (n = 3 per condition). (d) Log₁₀-transformed normalized expression of selected genes associated with ECM response (JAM2, PECAM1, PRELP), osteoclast-associated signaling and differentiation (TNFSF11/RANKL, ACP5/TRAP, CTSK), and inflammatory response (IL1B, CXCL8, STAT1). Data represent the mean ± standard deviation from N=3 biological replicates per condition. Statistical significance for panel d was assessed using two-tailed unpaired Student’s t-tests (*p < 0.05, **p < 0.01, ****p < 0.0001).

Gene ontology analysis of the top 100 genes upregulated in MIN relative to NM + GF revealed enrichment of biological processes associated with cell–matrix adhesion, cell migration and motility, and extracellular matrix organization (**Figure 4b, Supplementary Figure S13d**). Additional enrichment of developmental and vascular-associated processes suggested broader changes in cellular organization and interactions within the mineralized microenvironment. Together, these findings indicate that osteoclastogenesis within the engineered mineralized niche is accompanied by transcriptional programs emphasizing cell–matrix interactions and migration, distinct from the inflammatory signaling associated with conventional RANKL and M-CSF stimulation. Taken together with the transcriptional program in **Figure 2a-c** and the RANKL/OPG shift in **Figure 3b**, these data indicate that the mineralized niche supports osteoclast differentiation accompanied by adhesion-, migration-, and matrix-associated transcriptional programs that are distinct from the inflammatory response associated with exogenous RANKL and M-CSF.

Focusing on a curated set of genes associated with cell adhesion, matrix remodeling, osteoclastogenesis, and inflammatory signaling (**Figure 4c-d**), we found that mineralized constructs exhibited significantly higher expression of adhesion- and matrix-associated genes, including JAM2, PECAM1, and PRELP, alongside TNFSF11 (RANKL) and CTSK, a key protease involved in bone resorption. In contrast, conventional RANKL and M-CSF supplementation was associated with significantly higher expression of inflammatory mediators, including IL1B, CXCL8, and STAT1, while ACP5 (TRAP) expression did not differ significantly between conditions. Together, the transcriptional and secretory analyses demonstrate that osteoclastogenesis within the mineralized niche is accompanied by molecular responses distinct from those induced by conventional RANKL and M-CSF stimulation. While the mineralized microenvironment favors programs associated with cell adhesion, migration, and matrix remodeling, conventional growth factor supplementation elicits a more prominent inflammatory and chemokine-associated response. These findings highlight the influence of the engineered bone microenvironment on the molecular programs accompanying osteoclast differentiation and provide a basis for investigating the functional consequences of niche-driven osteoclastogenesis. This finding points to new opportunities in the field of engineered models that have traditionally relied on supraphysiological supplementation of cells with exogenous growth factors to stimulate biological activity. Here we show that such supplementation is not necessary when functional multicellular crosstalk is reconstituted within an engineered mineralized microenvironment.

### 2.4. The mineralized microenvironment supports functional resorption and recapitulates anti-resorptive therapy responses

Next, to evaluate whether our platform supports the functional response of osteoclasts, we compared matrix degradation in mineralized constructs that were identical in composition, both containing the embedded matrix cells and surface osteoblasts, and differed only in the presence (+Ocl) or absence (−Ocl) of macrophage-derived osteoclasts. Bone resorption was assessed by quantifying pit areas using second harmonic generation (SHG) imaging and by visualizing mineral degradation with microCT (**Figure 5a-d** and **Supplementary Figure S14**). Matrix degradation increased markedly in +Ocl constructs relative to −Ocl controls, which contained osteoblasts but no osteoclasts, supporting an additional contribution from osteoclasts beyond baseline remodeling by resident bone cells. To our knowledge, this dynamic, quantifiable level of resorption has not previously been demonstrated using radiologic (µCT) methods in an organ-on-a-chip. We further evaluated the functionality of osteoclasts differentiated in the chip by comparing their bone-resorbing ability to that of conventionally differentiated osteoclasts. Osteoclasts retrieved from the chip were seeded onto calcium phosphate-coated resorption plates and assessed for pit-forming activity. Their resorptive capacity was comparable to that of osteoclasts differentiated using exogenous RANKL and M-CSF (**Supplementary Figure S15**), indicating that osteoclasts matured on-a-chip retain functional bone-resorptive capacity even after removal from the device.

**Figure 5.**
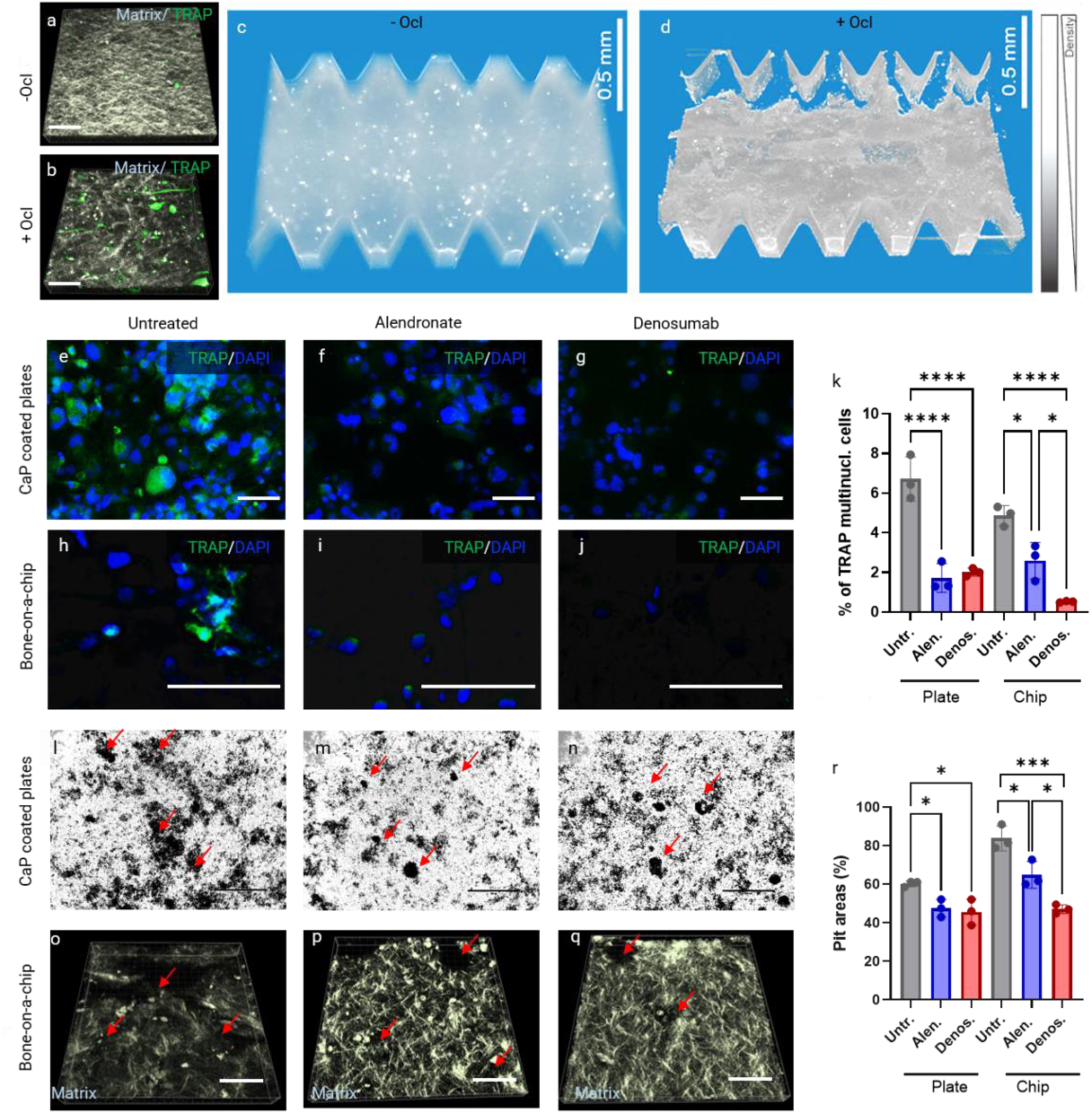
The mineralized bone-on-a-chip supports osteoclast-driven matrix resorption and anti-resorptive drug responses. Second harmonic generation (SHG) microscopy of mineralized constructs containing embedded osteoblast-lineage cells and surface osteoblasts, differing only in the presence (+Ocl) or absence (−Ocl) of macrophage-derived osteoclasts. The collagen matrix (SHG signal) was relatively preserved in the −Ocl condition (a), whereas the +Ocl condition (b) exhibited TRAP-positive cells (green) and localized loss of SHG signal, consistent with increased matrix degradation. Cross-sectional and lateral micro-computed tomography (micro-CT) reconstructions of −Ocl (c) and +Ocl (d) constructs further revealed structural alterations associated with osteoclast-mediated matrix resorption. Representative TRAP staining of multinucleated osteoclasts formed on calcium phosphate (CaP)-coated plates (e–g) or within the bone-on-a-chip platform (h–j) following treatment with vehicle, alendronate (10⁻⁷ M), or denosumab (30 µg mL⁻¹), showing reduced osteoclast formation following anti-resorptive treatment. Quantification of TRAP-positive multinucleated cells (k) confirmed suppression of osteoclast differentiation in both platforms. Representative resorption pits on CaP-coated plates (l–n) and localized matrix degradation visualized by SHG imaging on-chip (o–q), with red arrows indicating regions of resorption. Quantification of resorbed area (r) showed significant inhibition by both drugs, with denosumab producing a greater reduction than alendronate in the bone-on-a-chip platform under the conditions tested. Data represent mean ± SD from three biological replicates. Statistical significance was determined by one-way ANOVA followed by Tukey’s multiple comparisons test. For TRAP-positive multinucleated cells (k), all six groups were analyzed together, whereas resorption measurements (r) were analyzed separately for Plate and Chip conditions, with multiple comparisons performed within each platform. (*p < 0.05, **p < 0.01, ***p < 0.001, ****p < 0.0001). Scale bars: 100 µm (a, b, e–j, o–q, 400 µm (l–n) and 500 µm (c-d).

The inability of existing in-vitro models to capture key features of the bone microenvironment undermines their predictive power for evaluating anti-resorptive drugs. To evaluate whether our bone-on-a-chip system can resolve the differential activity of anti-resorptive drugs, we evaluated the effects of two clinically available anti-resorptive agents: alendronate, a bisphosphonate, and denosumab, a monoclonal antibody targeting RANKL. Both drugs are widely utilized in the treatment of osteoporosis and bone metastases, although clinical evidence consistently indicates denosumab’s superior efficacy^59^. Replicating these differential outcomes in-vitro has been challenging due to the distinct mechanisms of action of these drugs and their reliance on complex microenvironmental contexts. Denosumab inhibits osteoclastogenesis by binding to RANKL, thereby disrupting RANKL-driven osteoclast differentiation^60^. In contrast, alendronate is internalized by osteoclasts after binding to hydroxyapatite and interferes with cytoskeletal organization^61^. Importantly, these mechanisms depend on a mineralized matrix and the surrounding bone cells, an element largely absent in conventional in-vitro models, such as calcium phosphate-coated resorption plates. To capture this complexity, alendronate (10⁻⁷ M) or denosumab (30 μg/mL) was added to the culture medium of the mineralized constructs after the 3-day mineralization period, immediately before macrophages and surface osteoblasts were introduced. The construct was then cultured for 5 days. Cell viability remained comparable across untreated, alendronate-treated, and denosumab-treated chips after 5 days of treatment (**Supplementary Figure S16**). Samples were fixed and analyzed for cellular morphology, multinucleated cell formation, tartrate-resistant acid phosphatase (TRAP) expression, and functional osteoclast activity. Results were benchmarked against commercial calcium phosphate (CaP)-coated plates, which represent an industry-standard model of osteoclastogenesis using RANKL and M-CSF. Both drugs effectively suppressed osteoclastogenesis, as evidenced by a reduction in TRAP-positive multinucleated cells on both the chip and CaP plates (**Figure 5e-j, Supplementary Figure S17-19**). However, only on-chip assays assessing matrix resorption revealed a significant difference in drug efficacy that matches the reported clinical ranking of these drugs. Accordingly, on-chip experiments showed that denosumab significantly reduced bone resorption with smaller pit areas relative to both untreated controls and alendronate-treated samples. Because denosumab acts directly on RANKL, this result is consistent with the RANKL-dependent osteoclast activity that the mineralized niche promotes at the transcript and protein levels (**Figure 2b and Figure 3b**). Experiments off-chip did not recapitulate these clinically reported results (**Figure 5l-q**), highlighting the sensitivity of the bone-on-a-chip model in comparison to established methods.

### 2.5. Recapitulating features of tumor-bone interactions

To evaluate the ability of the bone-on-a-chip system to model complex, disease-relevant phenomena that depend on cross-communication between heterogeneous bone-resident cell populations, a challenge that remains difficult to address in-vitro, we investigated oral squamous cell carcinoma (OSCC) invasion into bone, a common and devastating process responsible for extensive bone destruction in oral cancer. Different from metastatic cancers, bone invasion occurs via cell infiltration into the matrix, as opposed to via vascular or lymph dissemination. While osteoclasts are known to facilitate bone invasion via paracrine signaling, existing in-vitro models fail to replicate the spatial and histological complexity observed in-vivo^62^. To address this gap, we compared OSCC invasion dynamics in engineered microenvironments with or without osteoclasts. Following osteoclast differentiation in the bone-on-a-chip system as described above, OSCC cells were seeded into one of the lateral channels of the microfluidic device after 3 days and co-cultured with bone cells (osteoblasts and pre-osteoclasts) for 24 h. Invasion was then evaluated in the presence or absence of osteoclasts. Fluorescently labeled (tdTomato) OSCC cells were used to visualize their spatial distribution relative to the mineralized matrix (**Figure 6a,g**), and pan-cytokeratin (PanCK) staining was used as a complementary epithelial marker (**Figure 6b,h**), given its reduced abundance in OSCC cells cultured on mineralized collagen (**Supplementary Figure S20**). We found significantly lower numbers of PanCK-positive cells in the lateral channels in the groups containing osteoclasts compared to those in groups without osteoclasts (**Figure 6b,h,p**). Consistent with these observations, analysis of tdTomato-positive cells revealed a greater number of migrating cells in the osteoclast-containing constructs (**Figure 6a,g,o)**. Meanwhile, in constructs where osteoclasts were not present, OSCC cells appeared largely localized along the mineralized matrix interface with limited penetration into the tissue (**Figure 6c,e,f**). In contrast, OSCC cells were detected beyond the bone-tumor interface, deeper into the regions between pillars of the device in osteoclast-containing samples (**Figure 6i,k,l**), These regions coincided with matrix disruption and cavity formation, as evidenced by localized loss of collagen signal in SHG imaging, while the collagen architecture remained relatively intact with limited penetration in the absence of osteoclasts (**Figure 6d,j**).

**Figure 6.**
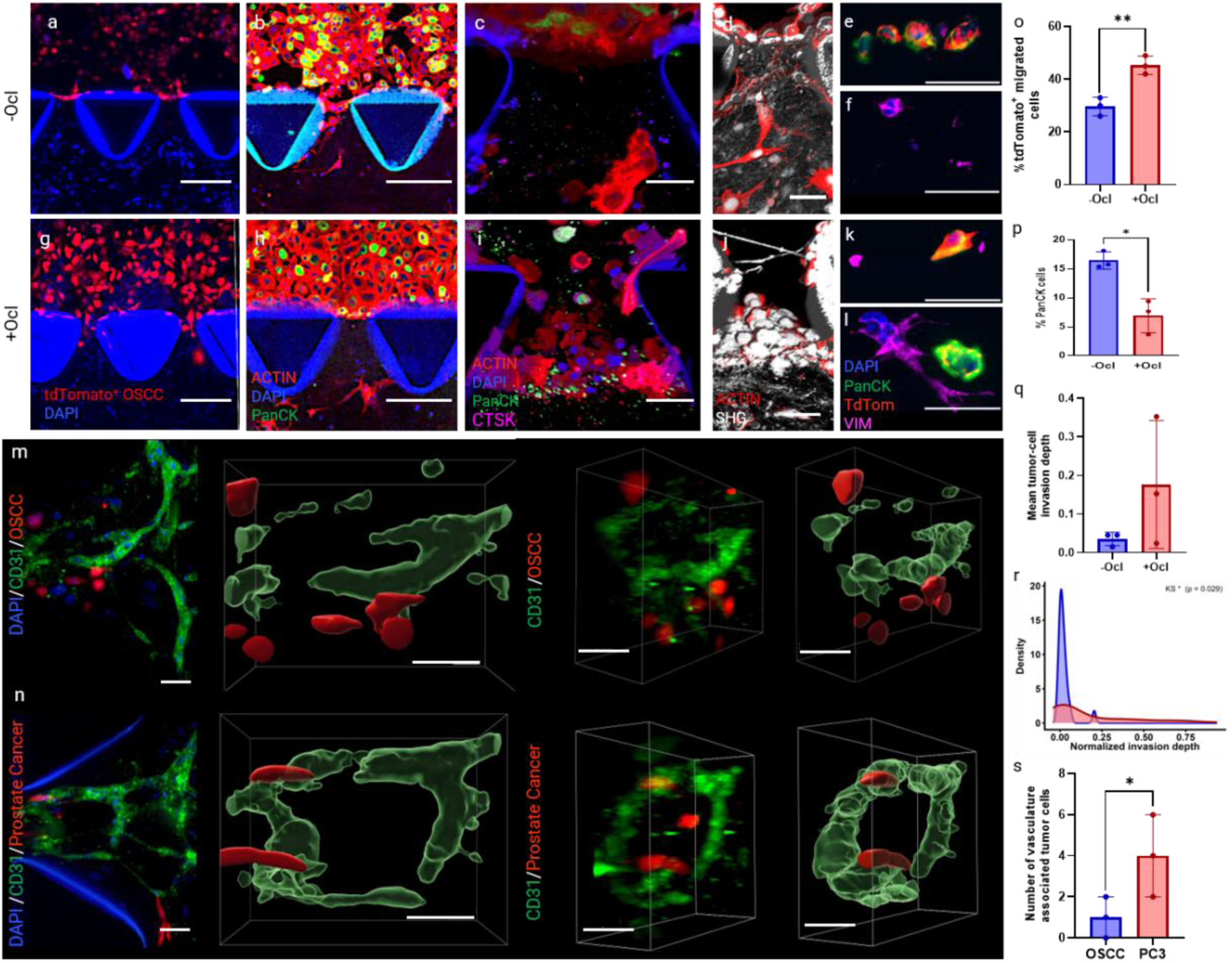
Bone-on-a-chip recapitulates key cellular and spatial hallmarks of OSCC bone invasion. Representative images of OSCC interactions with the mineralized bone microenvironment in the absence (a–f) or presence (g–l) of osteoclasts. (a, g) tdTomato-labeled OSCC cells (red) at the lateral channel–matrix interface, showing (o) significantly increased tumor cell migration in osteoclast-containing constructs.(b, h) Immunofluorescence staining for F-actin (red), DAPI (blue), and PanCK (green), showing (p) greater retention of PanCK-positive cells in the lateral channel in the absence of osteoclasts. Welch’s t-tests were used for (o, p) (n = 3 chips per condition). (c, i) Higher-magnification views of the tumor–matrix interface, stained for F-actin, DAPI, PanCK, and cathepsin K (CTSK; magenta), illustrating increased tumor cell penetration in osteoclast-containing constructs (d, j) Second harmonic generation (SHG; grayscale) and F-actin (red) imaging showing localized disruption of the collagen matrix and cavity formation in the presence of osteoclasts. (e, f, k, l) Immunofluorescence images of FFPE sections showing DAPI (blue), PanCK (green), tdTomato (tdTom; red), and vimentin (VIM; magenta), illustrating tumor cell localization and morphology relative to the mineralized matrix interface which corresponded to the (q) quantified mean normalized invasion depth of tdTomato-positive OSCC cells, calculated as the perpendicular distance from the matrix interface normalized to matrix thickness (n = 22 cells from 3 matrix-only chips; n = 58 cells from 3 osteoclast-containing chips). Mean invasion depth did not differ significantly between conditions (linear mixed-effects model with chip as a random effect, p = 0.27). (r) Distribution of individual normalized invasion depths, showing a rightward shift in osteoclast-containing constructs (Kolmogorov–Smirnov test, D = 0.35, p = 0.029) and increased variance (Levene’s test, F(1,78) = 7.64, p = 0.007). Representative fluorescence images and three-dimensional reconstructions of OSCC (m) and PC3 prostate cancer cells (n) within vascularized bone-on-a-chip constructs, showing tdTomato-positive tumor cells (red) relative to CD31-positive endothelial networks (green) and DAPI (blue). (s) Number of PC3 cells per chip exhibiting ≥50% overlap with the CD31-positive endothelial signal was significantly higher than OSCC showing greater vascular association of PC3 cells compared with OSCC cells (n = 3 chips per cancer type; Poisson regression, rate ratio = 4.0, p = 0.0317). Vascular overlap was used as an operational measure of association and does not independently establish intraluminal localization. Bar graphs demonstrate mean ± SD. Scale bars: (a, b, g, h) 200 µm; (c, d, i, j) 50 µm; (e, f, k, l) 50 µm; (m, n) 50 µm for fluorescence images, three-dimensional reconstructions, and magnified views. Statistical significance: *p < 0.05, **p < 0.01.

Immunofluorescence staining (PanCK/tdTom-OSCC/Vimentin/DAPI) further confirmed OSCC penetration into the mineralized matrix and demonstrated deeper, spatially heterogeneous tumor cell localization in the presence of osteoclasts (**Figure 6i-l**). To quantify invasion into the matrix, we measured the perpendicular distance of every tdTomato-positive tumor cell from the matrix interface and normalized this value to matrix thickness to account for construct variability (**Figure 6q**). Although the difference in mean invasion depth did not reach statistical significance (linear mixed-effects model with chip as a random effect, p = 0.27), distributional analysis revealed a significant rightward shift in invasion depth under osteoclast-containing conditions (Kolmogorov-Smirnov D=0.35, p=0.029) with increased variance in normalized invasion depth (Levene’s test, F(1,78) = 7.64, p = 0.007) and enrichment for deeply invading cells (18/58 vs 1/22 cells beyond 0.2 normalized depth, Fisher’s exact p = 0.017), indicating greater heterogeneity of invasion behavior (**Figure 6r**). Together, these findings suggest that osteoclast-associated matrix disruption may facilitate deeper penetration by a subset of OSCC cells, contributing to heterogeneous early invasion patterns within the mineralized microenvironment.

Since bone is targeted by tumors through fundamentally different routes, we next asked whether the platform could resolve these behaviors within a single engineered microenvironment. Whereas OSCC reaches bone by direct infiltration from an adjacent primary site, prostate carcinoma colonizes the skeleton following intravascular dissemination via hematogenous routes. To examine whether these distinct modes of engagement are reflected in our system, tdTomato-expressing OSCC and PC3 prostate carcinoma cells were introduced into vascularized bone-on-a-chip constructs containing osteoclasts, and their spatial association with CD31-positive vascular networks was quantified (**Figure 6m,n,s**). Tumor cells were classified as vasculature-associated using an overlap ratio threshold of 0.50 relative to the endothelial signal (CD31), without requiring definitive identification of intraluminal localization. Although tumor cells of both types were detected in proximity to vasculature, PC3 cells exhibited a significantly higher vascular association when compared to OSCC cells (**Figure 6s**). Three-dimensional rendering of representative images further illustrated these spatial localizations with examples of individual PC3 tumor cells localized within the vascular lumen (**Figure 6n**). Together, these preliminary observations demonstrate the ability of the platform to capture distinct patterns of tumor cell association with the vascularized bone microenvironment.

## 3. Discussion

Recent advances in bone tissue engineering, particularly in bone-on-a-chip platforms, have significantly improved our ability to model aspects of bone biology in-vitro^35, 63^. However, recreating a spatially organized, nanoscale-mineralized microenvironment that supports osteocyte-associated cellular responses and endogenous osteoclastogenesis, while accommodating vascular network formation without exogenous growth factors, remains a fundamental obstacle in skeletal in-vitro models^64^. To address this limitation, we demonstrate that engineering essential biomechanical and biochemical cues, specifically nanoscale mineralization and embedded osteocyte networks, drives spontaneous and functional osteoclastogenesis that mirrors hallmark features of bone remodeling. Moreover, the platform can be engineered to incorporate additional bone components, such as embedded vascular capillaries supported by stem cell-derived pericytes, enabling selective addition or removal of key cellular and matrix features that define bone function. In the absence of exogenous osteogenic factors, osteoblast-lineage cells embedded within a mineralized matrix adopted a mineral-responsive, osteocyte-associated state within three days, with elongated morphology, interconnected cellular networks, and higher number of SOST expressing cells (**Figure 2**). Bulk RNA-seq showed that this state was associated with enrichment of ossification, cytoskeletal organization, and matrix remodeling programs, whereas canonical osteocyte-associated transcripts did not exhibit coordinated upregulation, consistent with a partial shift toward an osteocyte-like phenotype without evidence of terminal maturation. This response is a step beyond existing 3D or on-chip bone models, which typically do not combine a nanoscale mineralized matrix with resident cells that respond to it. PDPN, a transmembrane glycoprotein, is typically induced when osteoblasts are subjected to mechanical stimuli within a mineralized environment and plays a crucial role in establishing the osteocytic network by mediating dendritic connectivity^65^. Importantly, both mineralized and non-mineralized constructs were maintained under identical rocking conditions, and computational modeling predicted minimal interstitial shear stresses, supporting matrix mineralization rather than rocking-induced flow as the primary contributor to the observed cellular responses. Although computational modeling predicted low interstitial shear stresses (∼0.001 Pa), a contribution from low-level oscillatory flow to osteocyte-associated marker expression cannot be excluded. Together with three-dimensional matrix encapsulation and cell–matrix interactions, these shared culture conditions may contribute to the osteocyte-associated features observed in NM constructs, while nanoscale mineralization promotes additional morphological and phenotypic changes. Collectively, these findings indicate that mineralization is associated with a shift in osteoblast-lineage cells toward an osteocyte-like, ossification-associated phenotype that is permissive for osteoclastogenesis, capturing essential features of bone physiology and remodeling in-vitro. The transcriptional program in mineralized chips was enriched in matrix remodeling, cytoskeletal organization, and bone-associated signaling, raising the possibility that nanoscale mineralization promotes an osteocyte-like phenotype through a cellular adaptation program that differs from conventional sequential osteoblast-to-osteocyte differentiation.

Our study establishes a self-regulating microenvironment that supports autonomous osteoclastogenesis^66^ mediated by multitypic cell-cell communication. The observed upregulation of RANKL and MMP9, alongside suppression of OPG, reflects intrinsic paracrine signaling between bone-forming and bone-resorbing cells, modulating osteoclast differentiation and activity. This signaling environment supports functional multinucleated osteoclasts and functional bone resorption, as validated by pit formation assays. Notably, macrophages differentiated into osteoclasts in the absence of exogenous RANKL/M-CSF supplementation during chip culture and localized near the mineralized matrix interface, suggesting effective niche recognition and activation. These findings are consistent with prior reports linking increased matrix stiffness to enhanced osteoclastogenesis, evidenced by the upregulation of canonical markers such as CTSK, MMP9, and ACP5^56, 67^. Neither the isolated matrix-associated cues nor conditioned medium reproduced the osteoclastogenic response observed in the complete mineralized construct under the conditions tested, suggesting that the combined cellular and matrix-associated environment is important for supporting osteoclast differentiation. However, in contrast to conventional in-vitro systems that rely on supraphysiological levels of RANKL and M-CSF, our platform leverages cues that arise from the mineralized multicellular niche to induce osteoclastogenesis. This physiologically relevant approach uncovered adhesion and junction-associated genes as potential regulators of osteoclast differentiation, features previously underappreciated^68^. Genes such as *FBLN5*, *CLDN3*, and *ADAM8* were significantly upregulated in the mineralized condition, suggesting a matrix-dependent transcriptional program. *FBLN5* was particularly noteworthy for its dual role in promoting *RANKL* expression while suppressing osteogenic signaling, potentially acting as a molecular switch favoring osteoclatogenesis^69^. *CLDN3*, a tight junction protein, may influence osteoclast-matrix interactions by modulating cell adhesion dynamics^70^. Elevated *ADAM8* expression, a known mediator of osteoclast precursor fusion, further suggests its potential involvement in osteoclast differentiation in the mineralized niche, since inhibition of *ADAM8* has been shown to markedly impair osteoclast formation^71^. By contrast, samples treated with RANKL/M-CSF exhibited a transcriptional profile characterized by higher expression of inflammatory mediators, including pronounced elevation of *IL1B* and *CXCL8*, in alignment with Luminex cytokine profiling (**Figure 3b** and **Figure S13c**). This divergence highlights distinct transcriptional signatures accompanying osteoclast differentiation under growth factor-supplemented and mineralized niche conditions. It is important to emphasize that this observation is based on a limited gene expression snapshot, highlighting the necessity of further validation across wider regulatory pathways.

The potential translational utility of our platform is illustrated by its ability to model therapeutic responses recapitulating clinically observed outcomes in-vitro. Clinical trials have demonstrated greater increases in bone mineral density with denosumab compared with alendronate^59^, yet recapitulating these effects in-vitro has proven difficult. Our system not only reflects these therapeutic trends but also establishes a robust preclinical framework for evaluating bone-targeted drugs. Importantly, most in-vitro models lack both osteocytes and the complete osteoblast-osteoclast-osteocyte axis of cellular crosstalk, often also excluding vascular or chemechanical cues modulating drug response, especially calcium and phosphate nanoscale minerals infiltrating collagens that cells bind and respond to. This omission limits mechanistic insight, particularly as osteocytes regulate bone remodeling through the RANKL/OPG signaling axis^72^. For example, the effects of alendronate on osteocyte biology remain poorly defined, despite its widespread clinical use^73^. Similarly, denosumab acts directly on RANKL to inhibit osteoclast differentiation and bone resorption^74^, highlighting the relevance of preserving endogenous RANKL signaling within engineered bone microenvironemnts. These findings represent an initial pharmacological proof of concept, as the experiments were conducted using single drug concentrations and a limited treatment window in a THP-1-derived model. Further dose-response studies, evaluation of different treatment stages, and validation using primary human cells will be necessary to establish the platform’s utility for predictive drug screening. By enabling integrated readouts across cell types within a physiologically mineralized biomimetic matrix, our platform provides a uniquely sensitive system for dissecting drug mechanisms, dose-response dynamics, and treatment withdrawal effects. Moreover, its modular design permits the incorporation of patient-derived cells, laying the groundwork for personalized medicine approaches in bone disease therapy.

Beyond its capacity to model resorption, our platform offers a unique opportunity to interrogate broader roles of osteoclasts in bone regeneration, osteocyte-mediated signaling, and cancer-bone interactions, including tumor invasion and metastasis^75, 76, 77^. One clinically significant application demonstrated here is the modeling of oral squamous cell carcinoma (OSCC) invasion, a process that targets mineralized tissues and contributes to catastrophic bone loss in oral cancer. Despite being the sixth most prevalent cancer worldwide^78, 79^, oral malignancies remain underrepresented in the development of advanced in-vitro models. Our system successfully recapitulates osteolytic behaviors characteristic of OSCC bone invasion, capturing for the first time in-vitro the spatial and cellular features observed clinically. This capability enables dynamic monitoring of tumor-bone interactions, which have historically been limited to static, end-point observations in-vivo. While prior studies have described bone invasion across multiple cancer types^62, 80, 81^, fundamental questions remain regarding the mechanisms driving OSCC tropism toward bone. For instance, epithelial-to-mesenchymal plasticity of OSCC cells at the tumor-bone interface has been proposed and may be triggered by biophysical and biochemical cues from the mineralized matrix^82, 83, 84^, highlighting the relevance of our system for dissecting the early steps of cancer cell adaptation to the bone microenvironment. Quantitative analysis of invasion depth revealed that osteoclasts induced a significant redistribution of invasion behavior, characterized by a rightward shift in the depth distribution and increased variance. This pattern suggests that osteoclast-mediated matrix remodeling does not uniformly enhance invasion across the tumor population but rather generates permissive microenvironments that enable deeper penetration by a subset of OSCC cells. Such heterogeneity mirrors clinical observations in which bone invasion often occurs through focal osteolytic niches rather than diffuse infiltration. Although osteoclast-associated remodeling preceded tumor seeding, these observations do not distinguish active matrix invasion from tumor cell migration into pre-existing permissive regions, and the ability of our platform to resolve these distributional changes highlights the importance of modeling spatially organized multicellular interactions within a mineralized context. The same principle applies to the comparison between distinct routes of skeletal engagement. Oral carcinoma reaches bone by contiguous infiltration from an adjacent primary site, whereas prostate carcinoma typically reaches bone through the circulation, and these trajectories place different demands on any model intended to capture them. That both behaviors were observed within our vascularized constructs, with prostate carcinoma cells enriched in vascular-associated positions relative to OSCC cells, indicates that the utility of the system is not restricted to local invasion. This comparison should be regarded as an initial demonstration rather than a mechanistic account, given the fact that the spatial metric employed does not resolve the sequence of events leading to vascular association.

Despite these advances, certain limitations remain. Raman analysis also revealed spatial variation in phosphate distribution across the engineered matrix, highlighting the need for future studies to examine how local differences in mineralization influence cellular behavior. The current model does not yet incorporate direct and controlled mechanical laoding, components of the adaptive immune system, such as T and B lymphocytes, which are known to influence both bone remodeling and tumor progression^85^. Additionally, while vascular and neural elements have previously been integrated into our mineralized matrix platform^38^, and the comparison presented here represents an initial application of the vascularized configuration to disease context, systematic incorporation of these features across the disease models will nonetheless require further optimization, particularly with respect to perfusion control and the temporal resolution of the tumor-endothelial interactions. Incorporating these components will expand the model’s utility for studying immune-mediated bone disorders and further enrich the complexity of the tumor microenvironment. Building on the observations reported here, the vascularized architecture of our model offers a promising foundation for investigating mechanisms of metastatic dissemination within a controlled, bone-like microenvironment. With its modular, plug-and-play design, this versatile platform is well-suited for studying bone biology across both physiological and pathological states, spanning development, homeostasis, and disease.

In summary, our biomimetic bone-on-a-chip model constitutes a significant advance in tissue-engineered New Approach Methodology (NAM) platforms for bone research. By recapitulating the structural, cellular, and biochemical intricacies of native bone, this system offers an unprecedented opportunity to investigate bone physiology, model disease progression, and evaluate therapeutic responses under physiologically relevant conditions. Its modular architecture and biological responsiveness position it as a versatile tool for both fundamental research and translational applications. As such, this innovation holds strong potential to accelerate the development of precision therapies and deepen our understanding of skeletal pathophysiology across diverse clinical contexts. More broadly, it marks a critical step toward the realization of high-complexity in-vitro human models capable of complementing animal models in the pursuit of mechanistic insights and human-relevant therapeutic discovery.

## 4. Methods

### Cell Culture

Human osteoblasts hFOB 1.19 (ATCC, CRL-3602) and human primary osteoblasts (Promocell, Inc, C12720) were cultivated in human osteoblast media (Cell Applications) supplemented with 10% fetal bovine serum (FBS) (Thermo Fisher) and 1% Penicillin/Streptomycin (P/S) (Thermo Fisher). The cells were used in passages 6-12. Human THP-1 monocyte cell lines (ATCC) were cultivated in Roswell Park Memorial Institute (RPMI 1640 media, Thermo Fisher) supplemented with 10% FBS and 1% P/S. Passages 10-15 were used to ensure consistent differentiation capacity. THP-1 cells were differentiated into macrophages by adding a solution of Phorbol 12-myristate 13-acetate (PMA, Sigma-Aldrich) (100 ng/mL) for 48h. The prostate cancer cell line PC3 (ATCC, CRL -1435) was obtained from Dr. Ryan Gordon (Knight Cancer Institute, OHSU). Human umbilical vein endothelial cells (Lonza) were cultured in Vasculife VEGF Endothelial complete media (EGM, Lifeline Cell Technology CA). Human mesenchymal stem cells (hMSCs; RoosterBio) were maintained in α-minimum essential medium (Gibco, Carlsbad, CA) supplemented with 10% FBS and 1% P/S. Experiments utilized HUVECs and hMSCs between passages 4 and 6. OSCC cells UCSF-OT-1109 (ATCC) were cultured in RPMI media supplemented with 10% FBS and 1% P/S. Osteoblasts were transduced with *pEF1α-tdTomato* lentiviral particles (Takara Bio) at a multiplicity of infection (MOI) of 5. Following transduction, cells were selected with puromycin (1 µg/mL) to establish a stable tdTomato-expressing population for visualization in Figure 1. UCSF-OT-1109 OSCC and PC3 cells were transduced with pEF1α-tdTomato lentiviral particles (Takara Bio) at a multiplicity of infection (MOI) of 5, and stable tdTomato-expressing cells were selected with puromycin (1 µg/mL) to visualize cellular invasion. All cells were maintained at 37 °C in a humidified incubator with 5% CO_2._

### Microfluidic device and osteoblast-laden gel

To establish the biomimetic tissue on a chip, we used a DAX-1 microfluidic device (AIM Biotech). Each device contains a main channel (1.30 mm wide x 0.25 mm high) flanked by two secondary media channels (0.5 mm wide x 0.25 mm high). Prior to gel loading, chips were coated with 1 mg/mL of poly-D-lysine (PDL; Merck) for three hours at 37°C to improve collagen adherence. To prepare the osteoblast-laden hydrogel, acid-solubilized type I collagen from rat tail (3 mg/mL, Thermo Fisher) was diluted to a final concentration of 2.5 mg/mL in 10x phosphate-buffered saline (PBS) with α-MEM media (Gibco, Thermo Fisher), neutralized with 1 N sodium hydroxide (NaOH) to pH 7.4. The mixture was combined with a suspension of osteoblasts for a final concentration of 3 × 10⁶ cells/mL. A 10 μL volume of the resulting pre-gel solution was pipetted into the central channel of each microfluidic device. The samples were maintained in a humidified incubator at 37 °C for 45 min to allow self-assembly fibrillogenesis.

For vascularized constructs, a suspension containing HUVECs, hMSCs, and osteoblasts in a 4:1:1 ratio (6 × 10⁶ HUVECs/mL, 1.5 × 10⁶ hMSCs/mL, and 1.5 × 10⁶ osteoblasts/mL) was dispensed into the main channel of the chip (10 μL total volume). The cell mixture was embedded in a hydrogel composed of 1.25 mg/mL collagen type I, 2U/mL thrombin, and 1.25 mg/mL fibrinogen. The vessels were allowed to form for 2 days in EGM prior to mineralization.

Assembly of the multicellular construct proceeded in two stages. First, osteoblasts embedded in the central-channel collagen (3 × 10⁶ cells/mL) underwent 3 days of matrix mineralization (see below), during which the embedded population differentiated toward an osteocytic phenotype within the mineralized matrix. After mineralization, a second population of osteoblasts (1 × 10⁴ cells) and THP-1–derived macrophages (3 × 10⁴ cells) were seeded into the lateral channels, yielding a construct containing three populations: osteocytes differentiated within the mineralized matrix, undifferentiated osteoblasts on the matrix surface, and macrophages that subsequently differentiated into osteoclasts. Upon addition of the second cell populations, the culture medium was switched to a 1:1 mixture of osteoblast medium (Cell Applications) and RPMI-1640, each supplemented with 10% FBS and 1% P/S. Unless otherwise stated, culture durations for osteoclastogenesis are reported from the time of macrophage addition (following the 3-day mineralization period).

### Nanoscale mineralization

To induce collagen mineralization, a mineralizing medium was prepared by mixing equal volumes of 9 mM CaCl_2·_2H2O (J.T. Baker) and 4.2 mM K_2H_PO_4 (_J.T. Baker) in α-MEM supplemented with 10% FBS and 1% P/S, based on our previously described protocol ^38^. Lacprodan^®^ OPN-10 (Arla Foods Ingredients Group P/S), a bovine milk-derived osteopontin analogue, was added to the CaCl_2-_containing solution at a concentration of 100 µg/mL before the addition of K_2H_PO_4._ To maintain a stable pH of 7.4, 25 mM HEPES was added to the mineralizing medium. The media was added to the side reservoirs of the microfluidic chips, and the devices were placed on a 2D rocker platform inside a humidified incubator (37 °C, 5% CO_2,_ 95% humidity) to allow continuous oscillatory flow and mineralization throughout the collagen matrix. Non-mineralized control chips and OIM-treated (osteoinductive medium) controls were maintained on the same 2D rocker platform for identical durations, ensuring that all groups experienced identical externally imposed rocking conditions. The media was replaced daily for 3 consecutive days to allow matrix mineralization of the biomimetic bone-on-a-chip construct.

### Matrix characterization

*SEM **-*** Mineralized and non-mineralized samples were fixed with 2% glutaraldehyde on a 2D rocker at room temperature for 2 hours, followed by washing with distilled water. Samples were dehydrated in an ascending ethanol series, removed from the chip, and subjected to critical point drying. The dried samples were mounted, sputter-coated with a 10 nm platinum coating, and imaged using a Helios Nanolab™ G3 DualBeam™ SEM (FEI, Thermo Fisher) (N = 3).

*FTIR -* Fourier Transform Infrared (FTIR) spectroscopy was performed using a Nicolet 6700 spectrometer (Thermo Scientific) in transmission mode. Spectra were recorded with 32 scans across a range of 4000–400 cm⁻¹, with a resolution of 4 cm⁻¹. The mineral-to-matrix ratio was determined by comparing the area of the ν₃PO₄ peak (1030 cm⁻¹) to that of the amide I peak (1660 cm⁻¹) after applying baseline correction and normalization to the amount of amide present. The crystallinity index was assessed using the splitting factor, calculated from the doublet peaks in the fingerprint region (500–650 cm⁻¹), primarily attributed to ʋ₄PO₄³⁻ bending vibrations. This calculation involved summing the peak heights at 565 cm⁻¹ and 605 cm⁻¹ and dividing by the height of the trough between them at 590 cm⁻¹. Peak height measurements were conducted using Spectragryph software, with spectra normalized to the intensity of the amide I band (1585–1720 cm⁻¹) following baseline correction (N=3).

*TEM –* For transmission electron microscopy, constructs were excised from the chips and fixed in 3% glutaraldehyde for 20 min at room temperature, then held in PBS. Samples were transferred to Dulbecco’s Modified Eagle Medium, post-fixed in 1% osmium tetroxide, and dehydrated through a graded ethanol series to 100%. Following a propylene oxide rinse, samples were infiltrated with Spurr’s epoxy and polymerized at 70 °C for 18 h. Ultrathin sections of 80 nm were collected and contrasted with saturated uranyl acetate followed by Reynolds’ lead citrate. Sections were imaged on an FEI G20 transmission electron microscope operated at 120 kV and recorded with an AMT XR41 2K × 2K camera (AMT, Woburn, MA). Constructs were sectioned perpendicular to the flow chamber, with one mineralized construct additionally sectioned parallel to the flow chamber. Mineralized specimens were examined at higher magnification to resolve intrafibrillar apatite crystallites within collagen fibrils (N = 3).

*Alizarin red -* An alizarin red assay was performed to confirm the presence of calcium deposition inside the hydrogel. The samples were stained with a 2% (w/v) alizarin red S solution, incubated for 15 minutes, and then washed in water until the solution was clear. The stained areas were analyzed by inverted microscopy (FL Autos, EVOS – Thermo Fisher Scientific) (N=3).

*Raman spectroscopy* - Samples were fixed in 4% paraformaldehyde for 20 min at room temperature. Raman spectra were acquired on a Horiba XploRA PLUS confocal Raman microscope (Horiba Scientific) using LabSpec 6 software for acquisition and cosmic-ray correction. The system was calibrated against a silicon wafer before each session. Samples were excited with a 785 nm laser at 285 mW, and signal was collected through a 600 gr/mm grating with a 10× objective over 200–2000 cm⁻¹. Constructs were kept on the chip to maintain consistent sampling relative to the reservoir and pillars. Eight points were collected along a transect from between the pillars and the stem, extending ∼200 µm toward the chip center and back, with the final point over plastic alone; chip-holder signal did not overlap the diagnostic bands at 958 and 1653 cm⁻¹. Spectra were baseline-subtracted, smoothed, and normalized to the amide I maximum (1653 cm⁻¹), except point 8, which was normalized to point 5 owing to plastic dominance over the pillars (N=3). For each construct, normalized spectra from seven spatial sampling locations within the collagen matrix (points 1–7) were averaged to obtain a representative spectrum for each biological replicate. These replicate-level spectra were subsequently averaged across independent constructs (MIN, n = 4; NM, n = 6) to generate the mean spectra for each condition. Point 8, corresponding to a plastic-dominated sampling region, was excluded from the matrix-averaged spectra and evaluated separately.

### Osteocyte characterization

#### Comparison of mineralized matrix and osteogenic induction

To compare the effects of the mineralized matrix with conventional osteogenic induction, collagen-containing osteoblasts (3 × 10⁶ cells/mL) were either cultured for 3 days to allow matrix mineralization or treated with osteogenic medium (100 nM dexamethasone, 50 μM ascorbic acid, and 10 mM β-glycerol phosphate) for 3 days. PDPN expression was used as an immunofluorescent marker to assess differences between conditions.

*Cell morphology and protein expression via immunostaining -* Samples (N = 3) were fixed in 10% neutral-buffered formalin for 30 min at room temperature and washed three times with PBS. All subsequent processing (including demineralization, permeabilization, blocking, antibody incubations, and washes) was performed in a microwave-assisted system using the PELCO Biowave Pro+ Tissue Processing System (Ted Pella, Redding, CA). Samples were placed on the PELCO ColdSpot® temperature-controlled surface within the PELCO EM Pro microwave vacuum chamber, sealed against the ColdSpot, and processed under a microwave power of 250 W with a vacuum of approximately 20 mm Hg. The ColdSpot® surface was maintained at 10 °C during demineralization, antibody infiltration, and wash steps, and at 21 °C during permeabilization and blocking. Demineralization was performed in 10% (w/v) EDTA in PBS (pH 7.4). Primary antibodies were then applied under microwave-assisted diffusion. The following primary antibodies were used: rabbit polyclonal anti-OCN (Bioss antibodies, bs4917R) (1:50 dilution), mouse monoclonal anti-PDPN (Origene, DM3500P) (1:100 dilution), DMP-1 (Thermo Fisher Scientific, PA5-57956) (1:100 dilution), FGF23 (Abcam, ab56326), (1:100 dilution), and SOST (Bioss bs-10200R 1:100 dilution). The following secondary antibodies were used at the specified dilutions: Alexa Fluor 555 goat anti-mouse IgG (Thermo Fisher Scientific, A21422) (1:200 dilution) and Alexa Fluor 647 goat anti-rabbit IgG (Thermo Fisher Scientific, A21244) (1:200 dilution). After washing, the secondary antibody (Alpaca anti-mouse Alexa fluor 488 antibodies, Jackson Laboratories, 615-545-214, 1:250 dilution) was applied using microwave-assisted diffusion. F-actin was stained with 555-conjugated phalloidin (molecular probes, Thermo Fisher), and nuclei were counterstained with DAPI (Thermo Fisher). Imaging was performed on a Zeiss LSM 880 confocal microscope, and quantitative analyses of cell morphology and protein expression were carried out with Imaris (v10.1, Oxford Instruments) and FIJI.

*Bulk RNA sequencing and transcriptomic analysis* **–** Osteoblasts encapsulated in collagen constructs (100 µL per construct) were cultured in 12-well plates in mineralizing and non-mineralized conditions as described above and collected after 3 days of culture. Four independent biological replicates were analyzed per condition. Total RNA was isolated from whole constructs using the Direct-zol RNA Microprep kit (Zymo Research). Because RNA isolated from the mineralized constructs showed greater degradation than RNA from non-mineralized controls, all samples were processed using an rRNA-depletion-based library preparation workflow to ensure consistent treatment across conditions. Libraries were sequenced using 150-bp paired-end reads to a depth of approximately 125–159 million read pairs per sample.

Raw sequencing reads were quality-filtered and adapter-trimmed using fastp (v1.4.0). Reads were aligned to the human GRCh38 reference genome using STAR (v2.7.11b) with gene annotations from GENCODE release 50. Gene-level read counts were obtained using STAR GeneCounts and independently assessed using featureCounts (Subread v2.1.1). Library strandedness was evaluated from strand-specific gene counts and was consistent with a reverse-stranded library configuration; reverse-strand counts were therefore used for downstream analyses.

Differential expression analysis was performed in R using DESeq2. Genes with fewer than 10 total reads across all samples were excluded prior to analysis. DESeq2 size-factor normalization was used to account for differences in sequencing depth, and differential expressions were calculated for MIN relative to NM samples. Positive log2 fold-change values therefore indicate higher expression in MIN constructs. Statistical significance was assessed using Benjamini–Hochberg-adjusted P-values, with a false-discovery rate (FDR) threshold of 0.05. Variance-stabilizing transformation (VST) was used for principal component analysis, sample-distance analysis, and expression heatmaps. For heatmaps, expression values were standardized independently for each gene as row-wise Z-scores.

Gene set enrichment analysis was performed using the **fgsea** R package^86^ on genes ranked by the DESeq2 Wald statistic. Hallmark gene sets and Gene Ontology Biological Process gene sets were evaluated, with positive normalized enrichment scores indicating enrichment in MIN and negative scores indicating enrichment in NM. Enrichment significance was assessed using Benjamini–Hochberg correction for multiple testing, with an adjusted *P* value (FDR) < 0.05 considered significant. Bone-related gene sets were subsequently examined to assess transcriptional programs associated with ossification, osteoblast differentiation, bone development, mineralization, cytoskeletal organization, and bone-remodeling functions. Selected gene-expression heatmaps were used to visualize bone signaling and osteocyte-associated functional genes, osteoblast and bone-matrix genes, and adaptive transcriptional responses across all biological replicates.

### Paracrine Signaling via Luminex

Secreted cytokines and growth factors were quantified using a ProcartaPlex multiplex immunoassay (Thermo Fisher), following the manufacturer’s instructions. Supernatants were collected from biomimetic bone-on-a-chip constructs after 1, 3, and 5 days of incubation under the following conditions: (i) mineralized chips, (ii) non-mineralized chips treated with RANKL (50 ng/mL, Peprotech) and M-CSF (30 ng/mL, Peprotech), and (iii) non-mineralized controls. The Luminex assay targeted 14 analytes associated with osteoclastogenesis: MCP-1 (CCL2), OPG, VEGF, IL-1β, GM-CSF, TNF-α, MIP-1β (CCL4), RANTES (CCL5), RANKL, M-CSF, MMP-9, GRO-α, MIP-1α (CCL3), and IL-6. Samples were run on a Luminex 200 instrument (Thermo Fisher). The analyte concentrations were determined by extrapolating individual experimental fluorescence intensity values against each analyte’s standard curve. The experiment was performed in three biological replicates, and the results were reported as individual concentrations (pg/mL) and as Z-scores for comparative heatmap visualization. For the groups treated with RANKL and M-CSF, the analysis was adjusted by subtracting the baseline concentrations already present in the medium.

### Osteoclast characterization

*THP-1 cells –*THP-1-derived macrophages were seeded into the lateral channels and cultured for 5 days under three conditions: (i) non-mineralized controls, (ii) non-mineralized chips supplemented with RANKL (50 ng/mL) and M-CSF (30 ng/mL), and (iii) mineralized chips. After culture, samples (N = 3) were washed with 1× PBS and fixed in 10% neutral-buffered formalin for 30 min at room temperature. All subsequent steps, including decalcification with EDTA, permeabilization, antibody incubations, and washes, were performed using a microwave-assisted system (PELCO Biowave Pro+, Ted Pella, Redding, CA). Samples were placed on the PELCO ColdSpot® temperature-controlled surface within the PELCO EM Pro microwave vacuum chamber, sealed against the ColdSpot, and processed under a microwave power of 250 W with a vacuum of approximately 20 mm Hg. The ColdSpot® temperature was maintained at 10 °C for antibody infiltration and washing steps, and at 21 °C during permeabilization. Permeabilization was performed with 0.1% Triton X-100, followed by three washes with PBS. Primary antibodies were applied under microwave-assisted diffusion. The following primary antibodies were used: anti-TRAP (Novus Biologicals, NBP2-45294, 1:100), anti-MMP9 (Bioss, bs-4593R, 1:100), and anti-cathepsin K (Thermo Fisher, PA5-18950, 1:100). After washing, secondary antibodies were applied using microwave-assisted diffusion: Donkey anti-goat Alexa Fluor 647 (705-607-003, 1:250 dilution), Donkey anti-mouse Alexa Fluor 790 (SA-000100, 1:250 dilution), and Alpaca anti-mouse Alexa Fluor 488 (615-545-214, 1:250 dilution).

*Patient samples -* All coded human subject samples, including peripheral blood samples and data, were obtained from the Cancer Early Detection Advanced Research (CEDAR) Specimen and Data Repository (IRB#18048). Samples under the repository are collected from clinics across OHSU following informed consent and released in accordance with the OHSU institutional review board. At the time of sample collection, donors were not undergoing any therapies or treatments. Peripheral blood monoclonal cells (PBMCs) were purified from whole blood over a Ficoll-Paque PLUS (GE Healthcare) gradient and cryopreserved prior to analysis.

*Monocyte isolation -* Monocytes were isolated using the Stemcell Technologies Human Monocyte Isolation Kit (Catalog# 19359). Briefly, PBMCs were thawed, washed and incubated with the isolation cocktail. After incubation with magnetic particles, the CD14+ cells were enriched using a magnet. CD14+ cells were counted and used for phenotyping and macrophage differentiation.

*Antibodies and flow cytometry* - On day 0 (prior to macrophage differentiation) and on day 7 (after macrophages were differentiated with M-CSF 30 ng/mL - Peprotech), a small aliquot of cells was stained to assess viability and cell surface marker expression. Cells were first incubated with a fixable live/dead dye to distinguish viable cells (BioLegend, Zombie Yellow Fixable Viability Kit, Catalog# 423104) and then stained with a combination of antibodies. Fluorescently labeled antibodies used were as follows: CD11b APC/Fire 750, CD14 APC, CD68 PE-Cy7, CD80 BUV805, CD86 BB515 and HLA-DR BUV563. Cell-surface staining was performed in FACS buffer (PBS supplemented with 1% FBS and 0.01% NaN3). Stained cells were acquired on the Cytek Aurora spectral flow cytometer. Data was analyzed with FlowJo software, version 10.10.0 (Treestar).

Cell morphology was visualized with Alexa Fluor 555-conjugated phalloidin, and nuclei were counterstained with DAPI (Molecular Probes, Thermo Fisher). Samples were imaged on a CrestOptics X-Light V3 spinning disk confocal system (Nikon Ti2) and a Zeiss LSM 880 laser-scanning confocal microscope (Fast Airyscan). Image analysis was performed with Imaris (v10.1, Oxford Instruments) and FIJI (ImageJ) to quantify multinucleated cells (≥3 nuclei per DAPI staining) and expression levels of TRAP, cathepsin K, and MMP9.

### Gene expression and quantification analysis

Samples were collected from three conditions: either non-mineralized, non-mineralized and treated with RANKL and M-CSF or mineralized. The lateral channels of each chip were washed with PBS and treated with TRI reagent (Zymo Research) for RNA/DNA/protein isolation. Total RNA was extracted and purified using the Direct-zol RNA MicroPrep kit (Zymo Research) following the manufacturer’s protocol. RNA concentration and purity were assessed using a Nanodrop One spectrophotometer (Thermo Fisher). Only samples yielding at least 150 ng of RNA were utilized for downstream analysis. A 1.5 μl aliquot of RNA lysate was used in the nanoString hybridization reaction, and the remainder was stored at −80 °C. nCounter Elements (nanoString) hybridization was performed according to the manufacturer’s instructions. The Pan-Cancer Progression panel was used, with the addition of a custom list of 55 bone-specific genes (see Dataset 1 for full gene list). Differential gene expression analysis was performed using the R package DESeq2 (v 1.24.0). Raw count data were normalized using DESeq2’s median of ratios method, and differential expression was assessed using the negative binomial generalized linear model (GLM). Genes with an adjusted p-value < 0.01 and absolute fold change ≥ 2 were considered significantly differentially expressed. Two differential gene lists were obtained by adjusting the p-value. Visualization was performed using the heatmap (v1.0.12) R package. A volcano plot was generated using glMDPlot and glXYPlot from the Glimma package (v1.10.1) to highlight differential expression between these two conditions.

### Bone remodeling in-vitro

*Calcium phosphate resorption assay -* Macrophages (THP-1 derived) were seeded at 1×10^5^ cells per well in a 48-well calcium phosphate (CaP)-coated plate (Bone resorption assay plate, AMSBIO) with RPMI media supplemented with 10% FBS and 1% P/S. The control groups for the experiment were: untreated macrophages (negative control) and a group treated with RANKL (50 ng/mL) and M-CSF (30 ng/mL) (positive control). To align the endpoints of the two arms, the CaP positive-control culture was started first; the chip cultures were initiated 4 days later, such that their 8-day protocol (3 days of mineralization followed by 5 days of osteoclastogenesis) concluded on day 12 of the control. At that point, cells harvested from the lateral channel of the biomimetic bone-on-a-chip were transferred onto the CaP substrate, and both groups were cultured together for a further 2 days (day 14) before analysis. The plates were then washed with PBS and deionized water, followed by 5 min of sodium hypochlorite, according to the manufacturer’s instructions. Subsequently, the samples were washed with DI water again and dried out before being taken to a microscope (Evos FL auto 2, Thermo Fisher). Pit areas were analyzed by ImageJ (images transformed into 8-bit and a threshold of 120 units) from 3 different biological replicates.

*Second harmonic generation microscopy -* Collagen matrix organization and integrity in mineralized chips were evaluated by second harmonic generation microscopy (SHG) in the presence (+OCl) or absence (-OCl) of macrophage-derived osteoclasts. Both conditions contained the embedded matrix cells and surface osteoblasts and differed only in the presence or absence of introduced macrophages that subsequently underwent osteoclast differentiation, allowing assessment of osteoclast-associated matrix remodeling relative to the resident cell baseline. The samples were fixed and immunostained for TRAP as previously described. The samples (N=3) were imaged using a laser-scanning confocal microscope (Zeiss LSM 880) configured to capture the reflected light between 485 nm and 495 nm, after exciting with a 900 nm laser. The 3D reconstruction was performed using Imaris 10.1 (maximum intensity projection, ch1 channel min-164 and max 6251.69, gamma 1.00), and the analysis of resorbed areas was performed by calculating the black (degraded tissue) to white (preserved tissue) ratio areas in FIJI (images transformed into 8-bit and a threshold of 30 units).

*Mineral visualization by X-ray tomography -* For assessment of mineralized regions of the bone-on-a-chip, the whole chip assemblies (n=3) were scanned via high-resolution X-ray tomography at 2.3 µm resolution (Zeiss XRadia 620 Versa X-ray Microscope). Chips were scanned after paraformaldehyde fixation (4%), rinsed in PBS, and sealed with Kapton tape. For visualization of the high-resolution X-ray tomography scans, samples were placed at distances from the X-ray source to produce the same energy intensity and resolution at the volume of interest. Scan settings were held constant across all samples as follows: 70 kV voltage, 8.5 W power, 4X objective, 2X binning, and 1-second exposure time. Images were transferred to Dragonfly Pro (Object Research Systems) for visualization. Intensity values varied between scans due to differing amounts of chip material remaining. For qualitative comparison, scans were window-leveled by adjusting the minimum intensity threshold to exclude the plastic chip from view while preserving all higher intensity values.

*Anti-resorption drug testing -* The most common anti-resorptive drugs, bisphosphonate alendronate sodium (USP Reference standard, Sigma Aldrich, 10^-7^ M) and the monoclonal anti-RANKL biosimilar Denosumab (Research Grade, Ichorbio, 30 µg/mL) were chosen for our study. Alendronate sodium was diluted in deionized water, and denosumab was diluted in PBS. Following mineralization, each drug was added to the culture medium and introduced into the lateral channels before macrophage and osteoblast seeding. Alendronate was used at 10⁻⁷ M and denosumab at 30 µg/mL. The media was changed daily for 5 consecutive days, after which cellular viability was evaluated for both drug conditions. The anti-osteoclastogenic effect of the drugs was determined by counting the number of TRAP^+^ and multinucleated cells in comparison with the untreated mineralized group. Osteoclast inhibition by both drugs was determined by quantifying the extent of resorption in FIJI (ImageJ), defined as the ratio of degraded (black pixels) to preserved (white pixels) tissue areas. The drugs were also tested on Ca-P coated plates (in-vitro gold standard for osteoclast activity) at the same concentrations to compare their osteoclast inhibitory effects for comparison with our biomimetic model.

### OSCC bone invasion

*2D experiments –* UCSF-OT-1109 OSCC cells were cultured in RPMI supplemented with 10% fetal bovine serum (FBS) and 1% penicillin/streptomycin (P/S) and used between passages 5–12. Type I collagen disks (100 μL per disk; 12 mm diameter) were prepared in 24-well plates and mineralized for 3 days using the mineralization medium described above. After mineralization, 5 × 10⁴ OSCC cells were seeded onto mineralized collagen, non-mineralized collagen, or tissue culture plastic controls and maintained for 3 days. Samples were then fixed with 4% paraformaldehyde and processed for immunofluorescence staining using mouse anti–pan-cytokeratin (Origene, TA190320; 1:1000) and rabbit anti-vimentin primary antibodies (NBP1-31327), followed by alpaca anti-mouse Alexa Fluor 488 (615-545-214; 1:250) and goat anti-rabbit Alexa Fluor 647 (Thermo Fisher Scientific, A21244; 1:200) secondary antibodies. Fluorescent images were acquired using an EVOS FL Auto imaging system (Thermo Fisher Scientific). Experiments were performed in biological triplicate (N = 3).

*OSCC model* - For the OSCC bone invasion test, UCSF-OT-1109 cells were cultured in RPMI supplemented with 10% FBS and 1% P/S in passages 5-12. Cancer cells (a total of 3×10^4^ cells) were added into the lateral channel of the microfluidic chips that contained early osteoclasts (after 3 days of osteoclast differentiation). In this configuration, the embedded bone cells were unlabeled, allowing OSCC cells to be identified directly by tdTomato fluorescence independently of marker expression. The co-cultures were maintained for 24 h to allow paracrine and direct interactions between OSCC cells and bone cells. Samples were fixed and immunostained for markers for osteoclasts (cathepsin-K, Invitrogen PA5-18950 1:100) and cancer cells (pan-cytokeratin, Origene SKU TA190320, 1:1000) as previously described, with OSCC localization determined from the tdTomato signal. Because matrix remodeling and osteoclast activity could alter the lateral channel–matrix boundary before tumor cell introduction, tumor cell position relative to the original interface alone was not considered sufficient evidence of invasion. Cells were evaluated for their three-dimensional spatial association with the surrounding collagen matrix, and only tumor cells identified within the matrix were included in the analysis. Cells occupying exposed regions or cavities within the central compartment without clear matrix association were not considered invading cells.

*Histological staining:* Tissue was fixed with 4% PFA for 45 minutes, then extracted from the microfluidic device using 2% agarose. Samples were processed for standard formalin-fixed paraffin-embedding by dehydration through a series of graded ethanols, xylene, and infiltration with paraffin wax. Processed tissue was embedded in paraffin wax for microtomy and sectioned at 5 µm thickness onto SuperFrost Plus charged glass slides. Sections were dewaxed and rehydrated for Hematoxylin and Eosin staining by graded ethanol washes followed by incubation in hematoxylin, bluing reagent, clearing agent, and eosin before dehydration and coverslipping. Brightfield images were acquired with a Zeiss Axiocam 712 color camera attached to a 0.63x camera mount on a Zeiss Axio Observer 7 inverted microscope stand.

*Immunofluorescent staining:* Rehydrated FFPE sections of extracted microfluidic chip samples were stained for protein targets on neighboring sections to histology. Antigen retrieval was performed using 1µg/mL proteinase K (Fisher BP1700), followed by blocking with 10% goat serum and overnight staining with primary antibodies (rabbit anti-Vimentin (Novus Biologicals NBP1-31327, 1:100), mouse anti-PanCK (Origene TA190321, 1:100), goat anti-tdTomato (OriGene AB8181-200, 1:100)). On day 2, slides were incubated for 2 hours at room temperature with secondary antibodies (donkey anti-rabbit 555 (Invitrogen A31572, 1:1000), donkey anti-goat 647 (Jackson Immuno 705-605-003, 1:1000), donkey anti-mouse 790 (Thermo Fisher SA-000100, 1:1000). Slides were then counterstained with NucBlue Fixed Cell ReadyProbes DAPI Reagent (Invitrogen R37606) and mounted with Prolong Diamond Antifade Mountant (Invitrogen P36970). Fluorescent images were acquired using a CrestOptics X-Light V3 spinning disk confocal system (Nikon Ti2) at 40X magnification.

*Invasion depth quantification:* Invasion depth was quantified from single-plane confocal immunofluorescence images in ImageJ for chips with or without osteoclasts. Nuclei were segmented from the DAPI channel by threshold-based binarization following background subtraction and size filtering to exclude speckle artifacts, and each nucleus was assigned a marker profile by scoring tdTomato, pan-cytokeratin, and vimentin signal within a fixed perinuclear ring; tumor cells were defined as tdTomato-positive. The invasion interface was defined relative to the original matrix surface boundary. For each tdTomato-positive tumor cell, the perpendicular distance from the matrix surface to the nuclear centroid was calculated and normalized to total matrix thickness to account for construct-to-construct variation. Cell-level depths were aggregated within biological replicates (58 tumor cells across 3 osteoclast-containing chips; 22 across 3 OSCC-only chips) prior to statistical testing.

### Cancer cell invasion and metastasis in vascularized constructs

To compare the invasive behavior of oral squamous cell carcinoma (OSCC) and prostate cancer cells within a vascularized bone microenvironment, vascularized constructs were generated as described above. HUVECs, hMSCs, and osteoblasts were embedded in a collagen–fibrin hydrogel at a 4:1:1 ratio, corresponding to 6 × 10^6^ HUVECs/mL, 1.5 × 10^6^ hMSCs/mL, and 1.5 × 10^6^ osteoblasts/mL. The hydrogel contained 1.25 mg/mL type I collagen, 2 U/mL thrombin, and 1.25 mg/mL fibrinogen. Constructs were cultured in EGM for 2 days to allow vascular network formation, followed by 3 days of mineralization. THP-1-derived macrophages were subsequently introduced and differentiated toward an osteoclast-like phenotype as described above. Following vascular network formation and macrophage addition, 2.10^4^ tdTomato-labeled PC3 prostate cancer cells or tdTomato-labeled UCSF-OT-1109 OSCC cells were introduced into a lateral channel of the device. The systems were maintained for 3 days to allow interactions between the cancer cells and the vascularized bone microenvironment. Samples were then fixed in 10% formalin (Fisher Scientific) and immunostained for CD31 (Abcam, ab215911; 1:100) to label HUVECs and cathepsin K (Invitrogen, PA5-18950; 1:100) to show osteoclast differentiation and activity. Nuclei were stained with DAPI (Invitrogen, R37606). Cancer cell localization was determined directly from the tdTomato signal. Fluorescence images were acquired using a Nikon Crest spinning-disk confocal microscopy system with a 20× objective. The proximity of migrated cancer cells to vascular structures within the vascularized bone microenvironment, as well as their spatial overlap with the vessels, was quantified using three independent replicates per cancer cell type using Imaris (v. 10.1, Oxford Instruments) (n = 3).

### Statistical analysis

All experiments were performed with at least three independent biological replicates unless otherwise specified. Data are presented as mean ± standard deviation (SD). Statistical analyses were performed using GraphPad Prism 10 and R, as appropriate. Comparisons between two groups were performed using two-tailed, unpaired Student’s t-tests, or Welch’s t-test where indicated (Figure 6o and 6p), while comparisons involving more than two groups were performed using ordinary one-way analysis of variance (ANOVA) followed by Tukey’s multiple-comparisons test, unless otherwise specified. Statistical significance was defined as p < 0.05.

For the anti-resorptive drug-response experiments (Figure 5), the percentage of TRAP-positive multinucleated cells was analyzed using a single one-way ANOVA encompassing all six experimental groups across the CaP-coated plate and bone-on-a-chip platforms, followed by Tukey’s multiple-comparisons test. Resorption measurements were analyzed separately for each platform because resorption was assessed using different methods: pit area on CaP-coated plates and SHG-based quantification of matrix degradation on-chip. Separate one-way ANOVAs followed by Tukey’s multiple-comparisons tests were therefore performed across the three treatment conditions within each platform.

Transcriptomic analyses were performed in R. For bulk RNA-seq, differential gene expression was assessed using DESeq2 following exclusion of genes with fewer than 10 total counts summed across all samples. Statistical significance was determined using Benjamini–Hochberg-adjusted P values, with an FDR < 0.05 considered significant; no additional fold-change cutoff was applied. Preranked gene set enrichment analysis was performed using fgsea, with genes ranked by the DESeq2 Wald statistic. Enrichment significance was assessed using Benjamini–Hochberg-adjusted P values, with an FDR < 0.05 considered significant. NanoString differential gene expression was analyzed using DESeq2, with significance defined by an adjusted p-value < 0.01. Comparisons of selected normalized gene-expression measurements were performed using two-tailed, unpaired Student’s t-tests.

For FGF23-positive cell quantification in vascularized constructs, differences between conditions were assessed using a linear mixed-effects model with Kenward–Roger-adjusted degrees of freedom and Holm correction for multiple pairwise comparisons.

To assess differences in tumor cell invasion depth between conditions, a linear mixed-effects model was fitted with the presence or absence of osteoclasts as a fixed effect and biological replicate (chip) as a random intercept to account for individual-cell measurements nested within chips. Differences in the distributions of normalized invasion depth were evaluated using a two-sample Kolmogorov–Smirnov (KS) test, and differences in variance were assessed using Levene’s test (Brown–Forsythe modification). These distributional analyses were performed on pooled individual-cell measurements and considered exploratory because they did not account for within-chip clustering. Fisher’s exact test was used to assess enrichment of deeply invading cells (normalized invasion depth > 0.2). For the vascularized cancer model, the number of tumor cells exhibiting ≥50% spatial overlap with CD31-positive endothelial structures was compared between OSCC and PC3 using Poisson regression with a log link, with cancer cell type as the predictor and independent chips as the unit of observation. Results were expressed as rate ratios. Confidence intervals were reported at the 95% level.

## Acknowledgements

We acknowledge funding from the Cancer Early Detection Advanced Research (CEDAR) Center at Oregon Health & Science University (OHSU) Knight Cancer Institute. NIH/NCI/NIDCR funding: R01DE029553, R01DE035326, R01CA310177, K99DE033689, T90DE030859, R90DE031533, and the Friends of Doernbecher Grant Program at OHSU. We also acknowledge the donation of Lacprodan® OPN-10 from Negin Mokhtari. Lastly, we acknowledge the Light Microscopy Core at OHSU for support with the image analysis.

## Competing Interest

Bertassoni LE holds equity and patents associated with HuMarrow Inc., RegendoDent Inc., NexOss Biotechnologies Inc., companies specializing in advanced tissue engineering and organs on chip, and other IP associated with regeneration technologies. The other authors declared that there are no conflicts of interest.

## Data availability

The RNA-seq and NanoString data generated in this study will be deposited in the Gene Expression Omnibus (GEO) (accession number to be added) before publication. Source data are provided with this paper. Owing to their size, raw imaging data are available from the corresponding author upon request.

## Ethics declaration

All human samples and data were coded and obtained from the Cancer Early Detection Advanced Research (CEDAR) specimen and data repository at OHSU (IRB #18048). Samples in the repository are collected from clinics across OHSU following informed consent and are released in accordance with the OHSU Institutional Review Board. Primary osteoblasts and HUVECs were purchased from commercial vendors.

## Author Contributions

M.G.C.S. and A.A. contributed equally to this work. M.G.C.S. and A.A. conceived the study, designed and performed experiments, analysed data, prepared figures, and wrote the original draft of the manuscript. M.H. performed the oral squamous cell carcinoma and prostate cancer invasion experiments, analysed data, and helped prepare the figures. D.M.R. analysed the NanoString data, quantified cancer cell invasion depth and vimentin staining, and helped prepare the figures. G.E.R. acquired and analysed the micro-CT data. R.D. isolated the PBMCs and performed the flow cytometry. R.T. performed the computational simulations. D.B. acquired and analysed the confocal Raman data. M.A.A.F. performed osteoclast immunostaining and quantification. S.M.V. performed SEM imaging and analysis and the Luminex assays and analysis. A.D. performed the bioinformatic analysis of the NanoString data. J.L. performed RNA quantification and ran the NanoString assays. J.V.N. performed image quantification for the cancer invasion experiments. D.F.J. performed quantification for the paracrine signalling and cancer invasion experiments. A.S.P.L. acquired the micro-CT data. E.O. provided expertise in osteoclast biology and helped identify and analyse osteoclasts. C.M.F. performed osteoclast imaging, second harmonic generation imaging, and imaging of the drug treatment experiments. R.E.G. contributed to the conceptualisation of the study. L.E.B. conceived and supervised the study, acquired funding and revised the manuscript. All authors reviewed and edited the manuscript and approved the final version.

## Notes

### Competing Interest Statement

Bertassoni LE holds equity and patents associated with HuMarrow Inc., RegendoDent Inc., two companies specializing in advanced tissue engineering, and other IP associated with regeneration technologies. The other authors declared that there are no conflicts of interest.

### Summary of Updates

Figures 2, 3, and 6 were updated.

