## supporting information for "Endogenous Osteoblast-Osteocyte-Osteoclast Signaling Enables Bone Remodeling, Drug Response, and Cancer Invasion in a Nanoscale Calcified Bone-on-a-Chip"

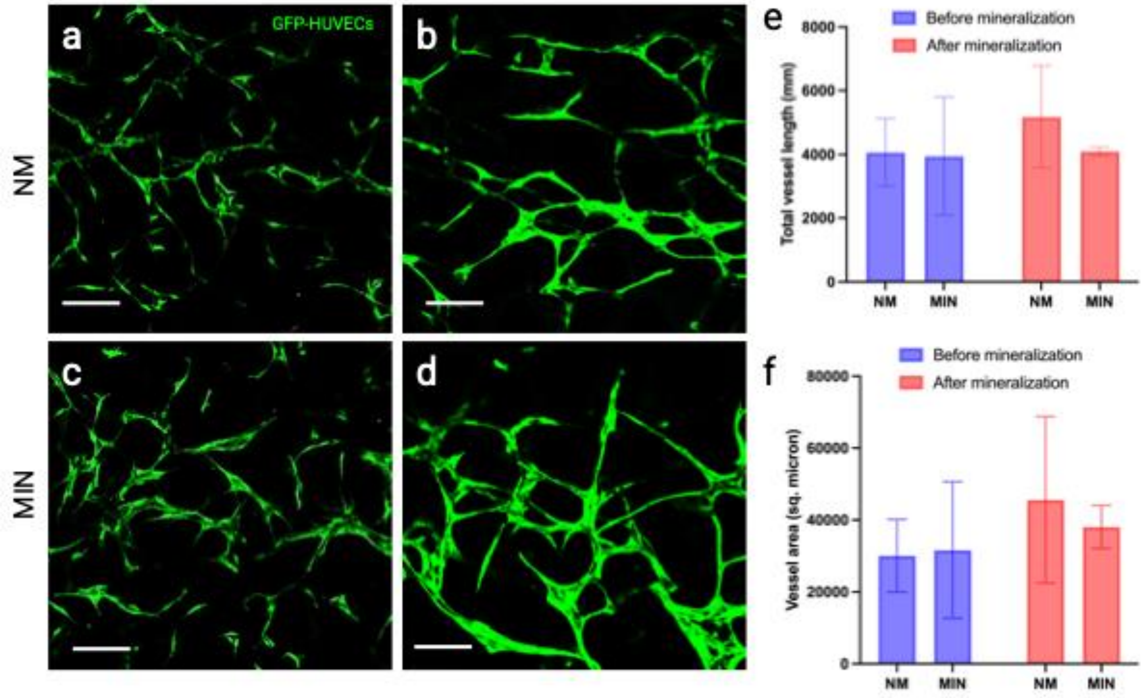


**Figure S1. Bone-on-a-chip supports the addition of vascular networks.** Representative images (a-d) showing comparable endothelial capillary network formation by GFP-expressing HUVECs (Angioproteomie) encapsulated in non-mineralized (a, b) and mineralized samples (c,d) on day 2 (a, c) and day 5 (b, d) of culture. Quantification of the total vessel length (e) and vessel area (f) showed no significant differences between mineralized and non-mineralized samples before or after mineralization. (n = 3 chips per condition) Bar graphs show mean ± SD. Scale bars: 30 µm (a,c) and 10 µm (b,d).

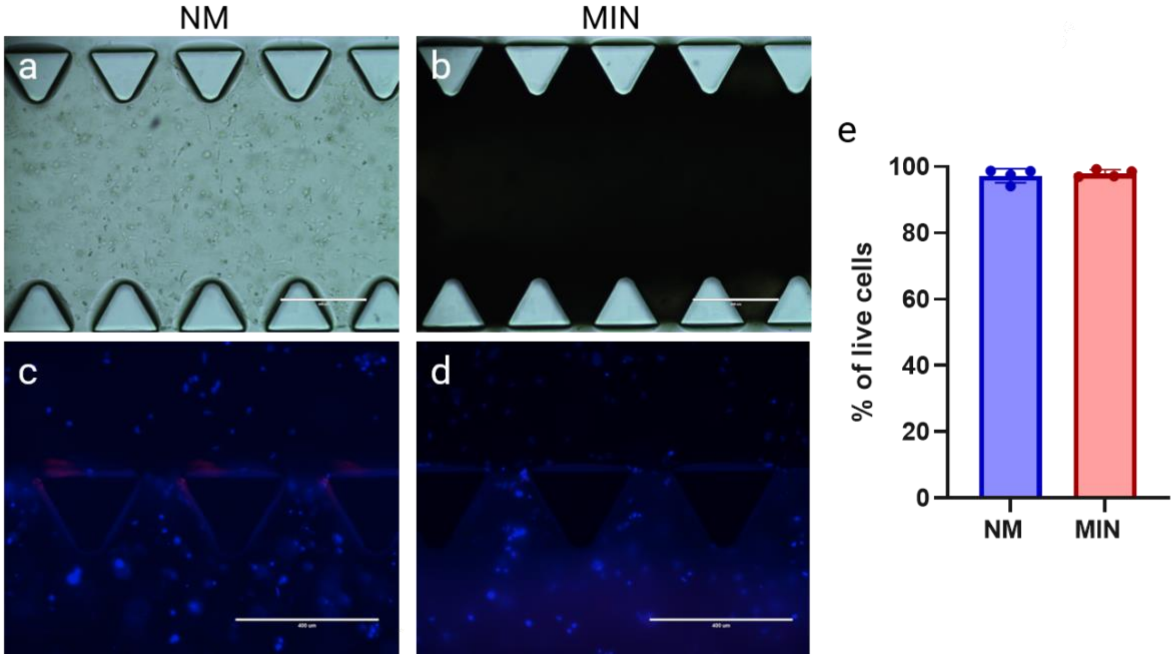


**Figure S2. Characterization of on-chip mineralization process and associated cellular viability.** Incorporation of calcium phosphate in the mineralized groups is evident from the change in opacity in phase contrast images (a,b) when compared to the non-mineralized samples. Representative images (c, d) of chips stained with Hoechst (blue) and propidium iodide (dead, red) following 5 days of culture in non-mineralized (NM) and mineralized (MIN) conditions showed minimal cell death in both groups, with (e) showing no statistically significant difference in viability between the groups after a t-test analysis (N=4). Bars represent mean ± SD. Scale bars in a-d represent 400 µm.

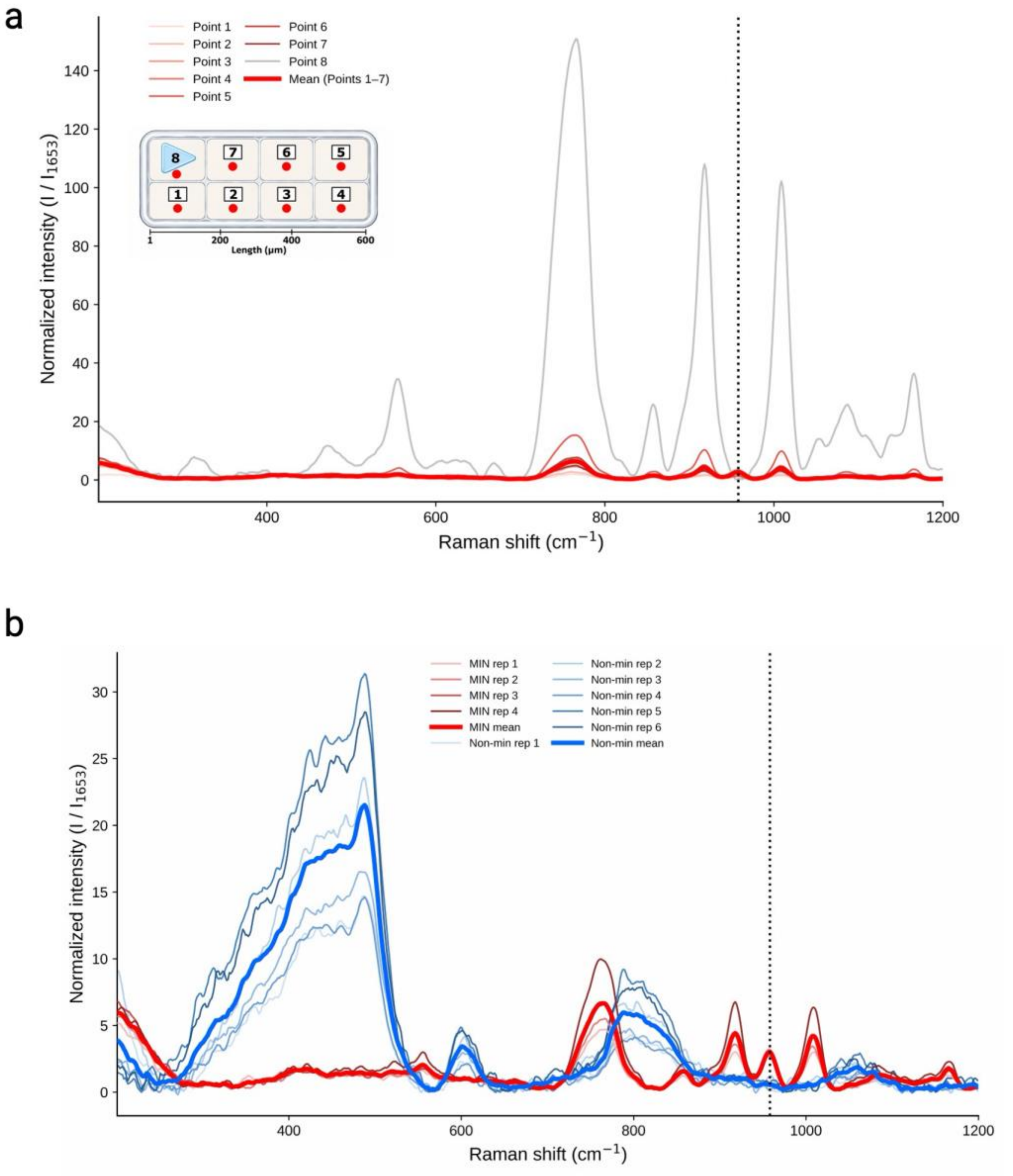


**Figure S3**. **Spatially resolved Raman characterization of mineralized and non-mineralized bone-on-a-chip constructs.** (a) Raman spectra acquired at eight positions across the chip, as indicated in the inset schematic. Points 1-7 represent locations within the collagen matrix, and point 8 corresponds to the device's pillar feature. The mean spectrum of points 1-7 is shown separately. (b) Raman spectra from individual mineralized (MIN) and non-mineralized (NM) replicates, averaged across the sampled matrix locations, with the corresponding group means highlighted. Spectral intensities were normalized to the amide I band (1653 $cm^{-1}$), and the dotted vertical line indicates the phosphate $\nu_1$ band (~960 $cm^{-1}$).

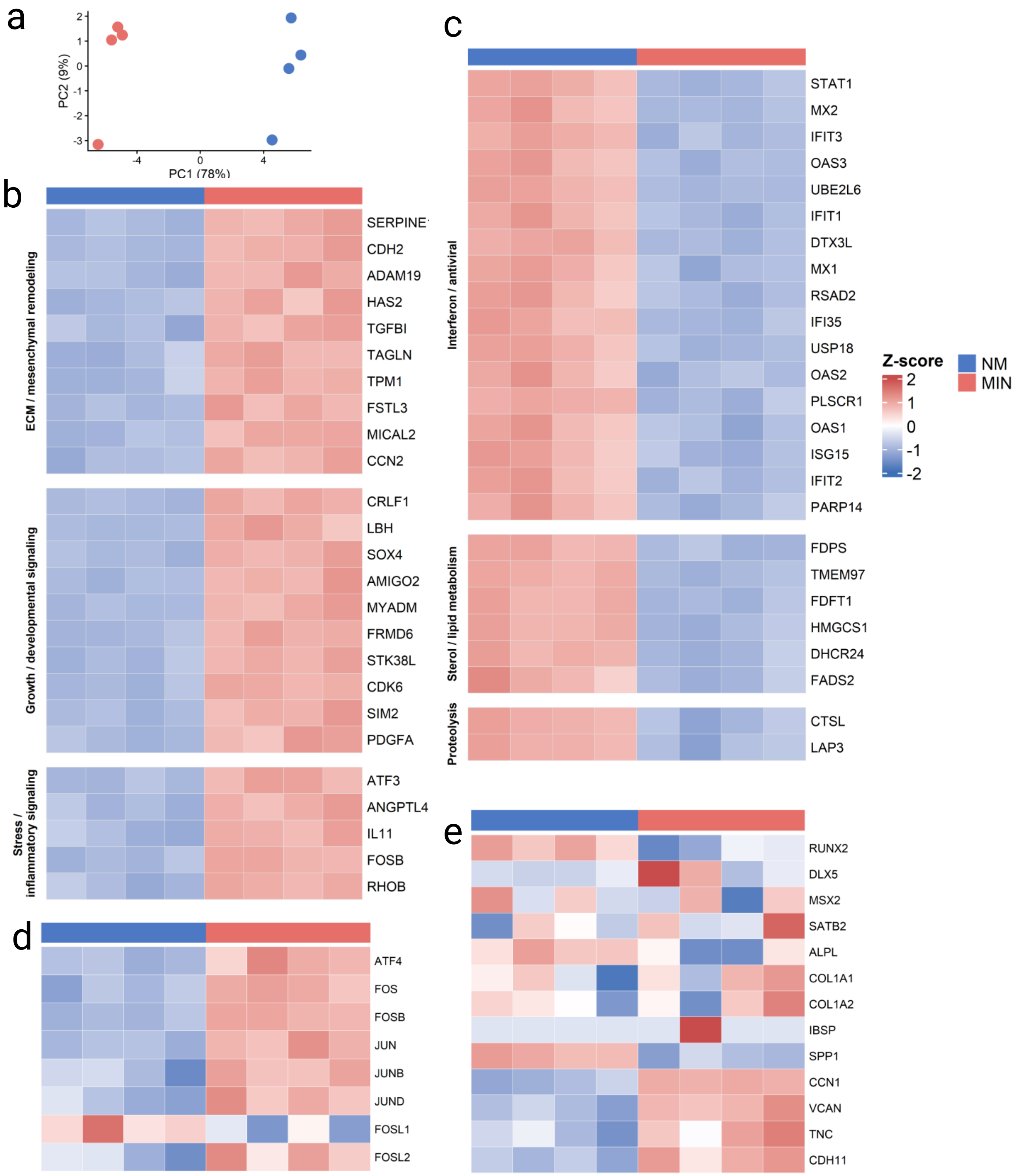

a
PC2 (9%)
PC1 (78%)
b
ECM / mesenchymal remodeling
SERPINE1
CDH2
ADAM19
HAS2
TGFBI
TAGLN
TPM1
FSTL3
MICAL2
CCN2
Growth / developmental signaling
CRLF1
LBH
SOX4
AMIGO2
MYADM
FRMD6
STK38L
CDK6
SIM2
PDGFA
Stress / inflammatory signaling
ATF3
ANGPTL4
IL11
FOSB
RHOB
c
Interferon / antiviral
STAT1
MX2
IFIT3
OAS3
UBE2L6
IFIT1
DTX3L
MX1
RSAD2
IFI35
USP18
OAS2
PLSCR1
OAS1
ISG15
IFIT2
PARP14
Sterol / lipid metabolism
FDPS
TMEM97
FDFT1
HMGCS1
DHCR24
FADS2
Proteolysis
CTSL
LAP3
Z-score
NM
MIN
d
ATF4
FOS
FOSB
JUN
JUNB
JUND
FOSL1
FOSL2
e
RUNX2
DLX5
MSX2
SATB2
ALPL
COL1A1
COL1A2
IBSP
SPP1
CCN1
VCAN
TNC
CDH11

**Figure S4. Transcriptional differences between non-mineralized and mineralized osteoblast-lineage cell cultures.** (a) Principal component analysis (PCA) of bulk RNA-seq samples from non-mineralized (NM) and mineralized (MIN) conditions showing separation primarily along PC1. (b) Heatmap of genes most strongly increased in MIN samples, including genes associated with extracellular matrix and membrane remodeling, growth and developmental processes, and stress-responsive transcriptional programs. (c) Heatmap of genes most strongly increased in NM samples, including genes associated with interferon/antiviral responses, sterol and lipid metabolism, and proteolysis. (d) Heatmap of selected genes associated with an adaptive/AP-1 transcriptional response. (e) Heatmap of selected osteoblast- and bone matrix-associated genes. Heatmaps display row-wise Z-scores of variance-stabilized expression values, with red indicating higher and blue indicating lower relative expression for each gene across samples. Each column represents an individual biological replicate.

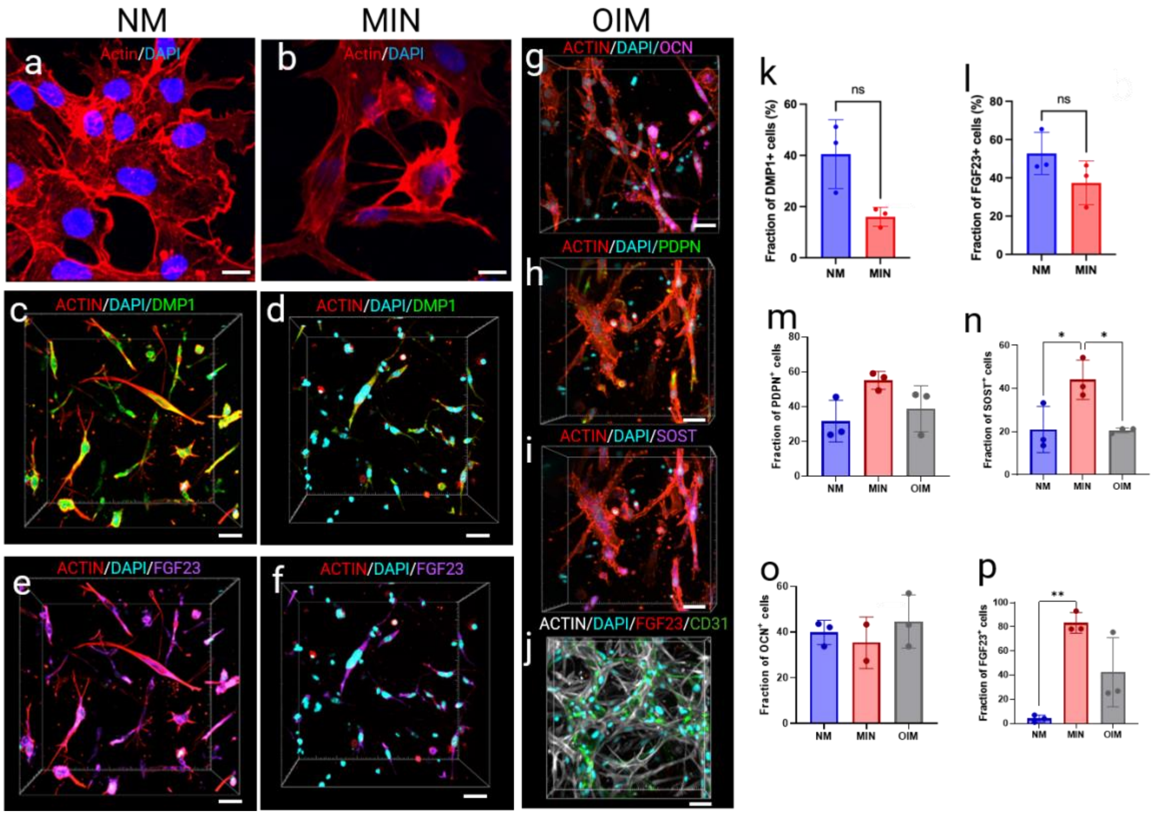


**Figure S5. Osteocyte-associated morphological changes and marker characterization in mineralized and non-mineralized constructs.** Representative confocal images (a,b) of samples stained for F-actin (red) and DAPI (blue) after 3 days in culture show distinct cellular morphologies between non-mineralized (NM) and mineralized (MIN) conditions, with MIN samples displaying dendritic cytoplasm extensions consistent with an osteocyte-like morphology. Representative images show DMP1 (green, c,d) and FGF23 (purple, e,f) immunostaining of osteoblast lineage cells cultured for 3 days under NM (c,e) and MIN conditions (d,f). Osteoblast associated markers OCN (magenta, g), PDPN (green, h) and SOST (purple, i) was also assessed in osteoblast lineage cells cultured for 3 days in osteoinductive media (OIM). Quantification shows the percentage of DMP1-positive (k) and FGF23-positive (l) cells in NM and MIN osteoblast cell-line constructs, and PDPN-positive (m), SOST-positive (n), and OCN-positive (o) cells across the indicated experimental conditions where SOST positivity was significantly higher in MIN than in NM and OIM. Separately, distinct from the osteoblast cell-line experiments shown in panels e, f, and l, FGF23 expression was evaluated in vascularized constructs incorporating primary human osteoblasts and cultured in OIM for 3 days (j), and FGF23 positivity was found to be significantly higher in MIN than in OIM (p). Statistical significance is indicated in the graphs (*$p < 0.05$; $p < 0.01$; ns, not significant). Bars represent mean ± SD. Scale bars: 20 µm (a-j).

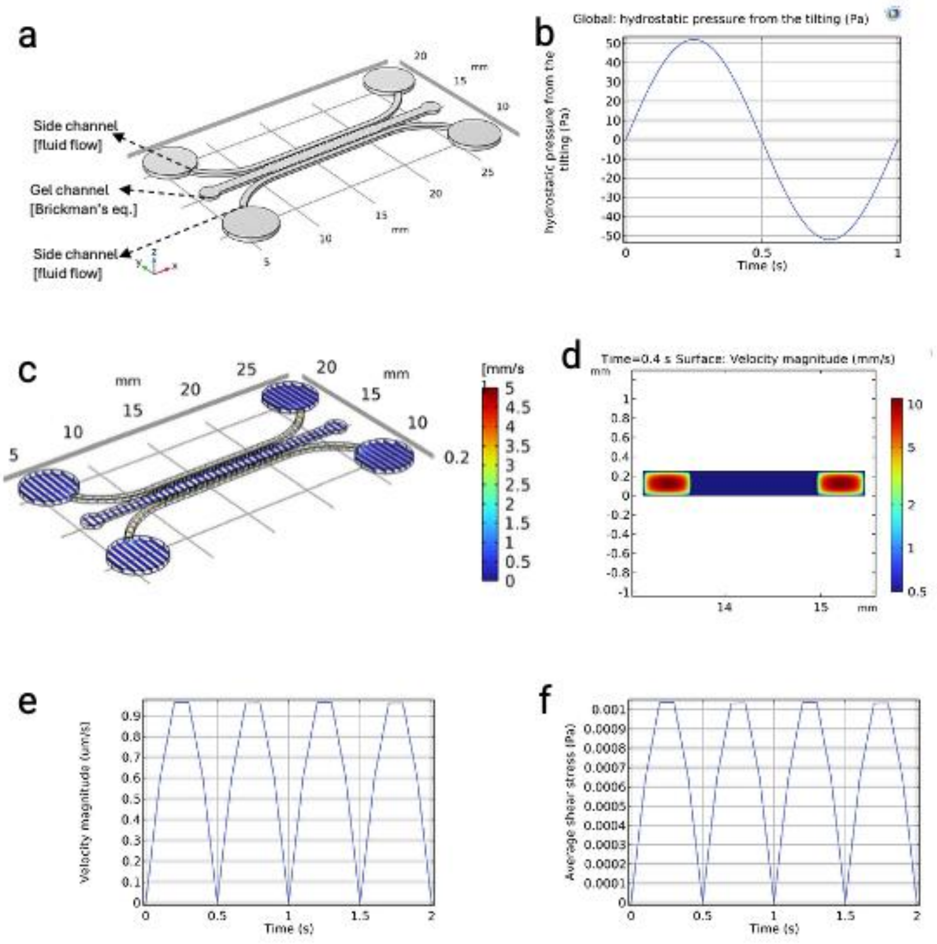


**Figure S6**. **Computational fluid dynamics (CFD) modelling of the bone chip**. (a) Computation domain was divided into a free-flow media domain to represent side-channels and a porous matrix domain that corresponds to collagen. (b) The plot shows a sinusoidal pressure boundary condition as a function of time at the chip inlet reservoirs to mathematically approximate the rocking boundary condition. (c) Velocity field plotted over the entire computational domain using several 2D slices and (d) an example yz-cross-sectional slice showing velocity heat map. Average velocity magnitude (e) and resulting shear stress (f) plotted over time in the porous matrix domain.

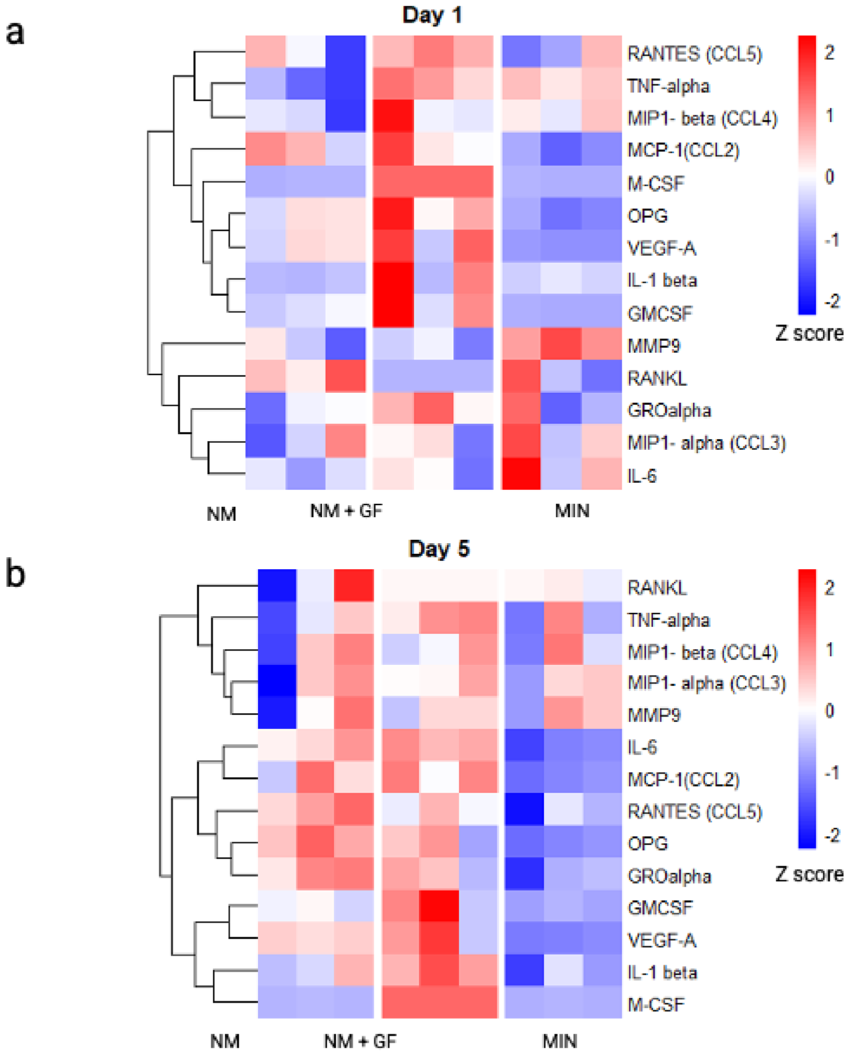


**Figure S7**. **Relative secretion profiles of osteoclastogenic cytokines in mineralized and non-mineralized bone-on-a-chip cultures.** Heatmaps showing relative secretion of 14 proteins/cytokines and chemokines in the supernatants of non-mineralized (NM), non-mineralized + RANKL/M-CSF (NM + GF), and mineralized (MIN) samples after 1 (a) or 5 days (b) of culture as measured by Luminex multiplex analysis. The results are presented as Z-scores, with measurements obtained from 3 different biological replicates. Exogenously supplemented RANKL and M-CSF concentrations were subtracted from the corresponding NM + GF measurements before analysis.

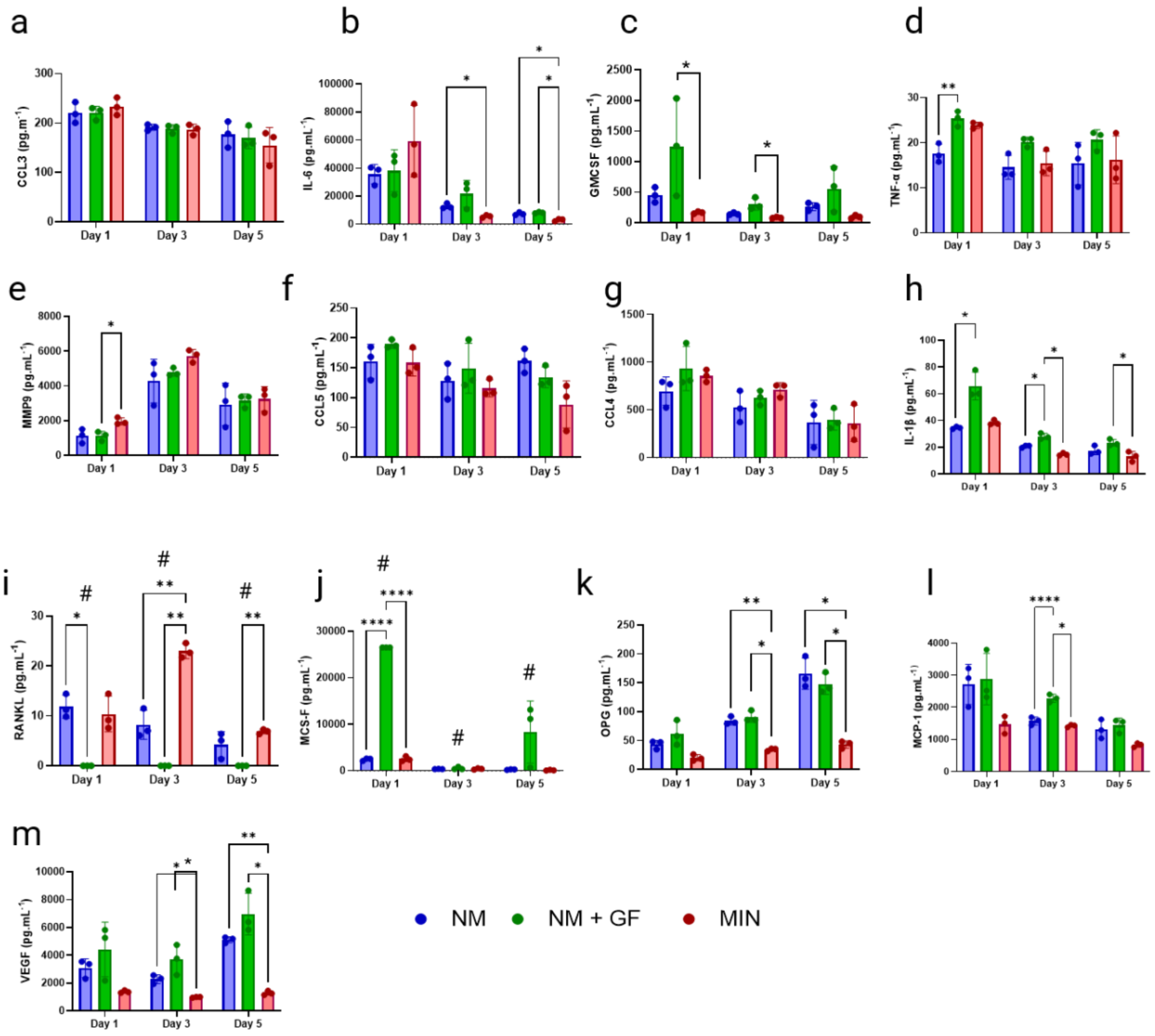


**Figure S8**. **Proteins/cytokines and chemokines related to osteoclastogenesis in solution after 1, 3, or 5 days of culture, as measured by Luminex multiplex analysis.** The average measured concentrations of proteins/cytokines (N=3) are represented in pg/mL. Significant statistical differences are represented by * $p<0.05$, ** $p<0.01$, *** $p<0.001$, and $p<0.0001$ after one-way ANOVA with Tukey's correction. # represents groups that already had RANKL (50 ng.mL$^{-1}$) and M-CSF (30 ng.mL$^{-1}$) in the media (NM + GF group in panels i and j). These known concentrations in the media were subtracted from the measured Luminex values in those plots. Bars represent mean ± SD.

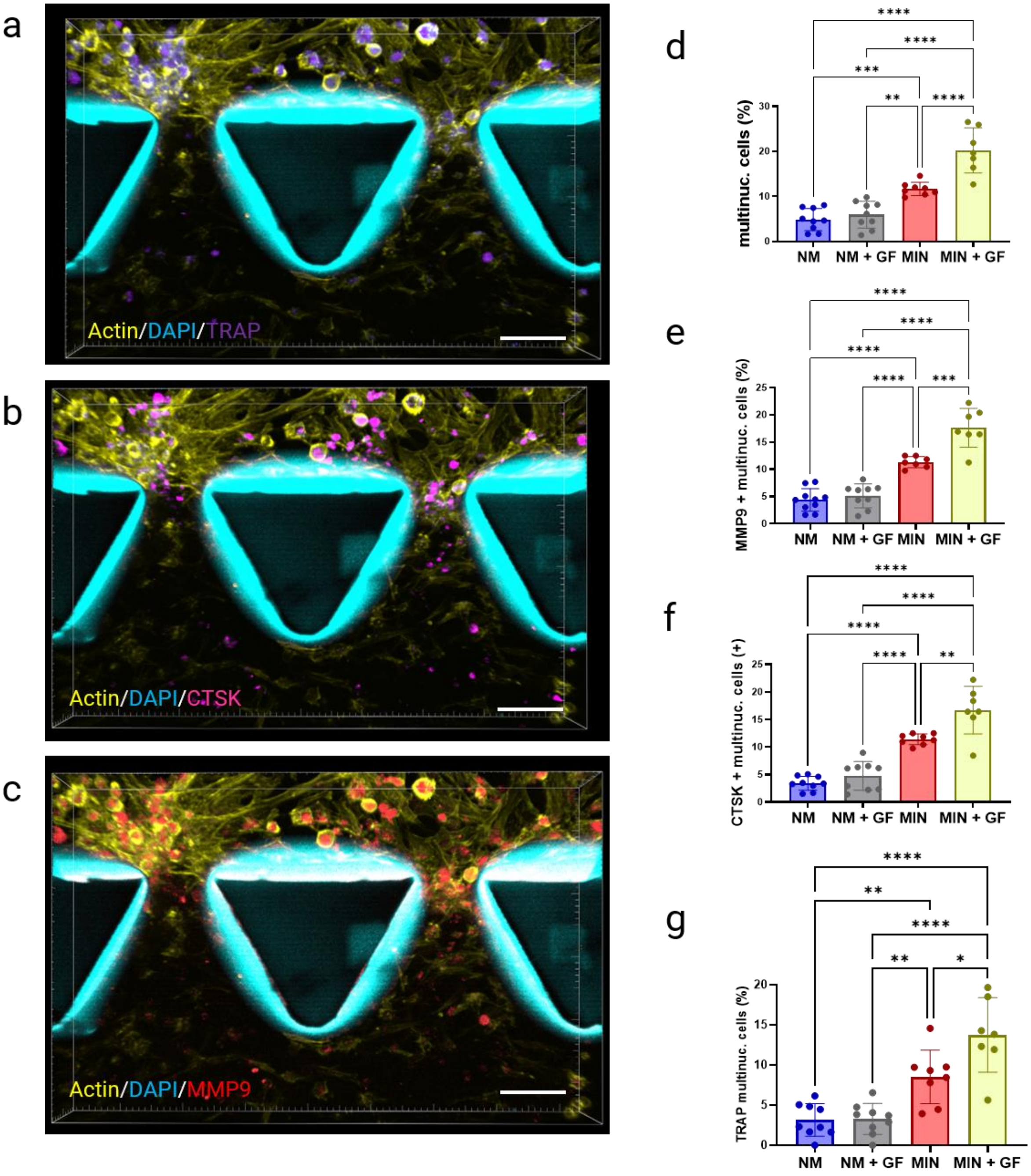


**Figure S9**. **Osteoclast differentiation in the bone-on-a-chip is enhanced by mineralization and further enhanced by RANKL/M-CSF.** Representative confocal images (a-c) of mineralized samples treated with RANKL (50 ng/ml) and M-CSF (30 ng/ml) supplemented medium show cells positive for TRAP, CTSK, and MMP9, respectively. Graphs (d–g) illustrate the increase in fractions (n = 3); data are mean ± standard deviation) of multinucleated cells (d) and MMP9-positive (e), CTSK-positive (f), and TRAP-positive (g) multinucleated cells depending on mineralization and the addition of growth factors (GF) (N=3). Mineralized samples supplemented with GF exhibited the highest fractions of multinucleated cells and MMP9-positive, CTSK-positive, and TRAP-positive multinucleated cells, indicating that RANKL and M-CSF further promote osteoclastogenesis of macrophage-derived progenitors under mineralized conditions.

Statistical significance in (d–g) is indicated as *p<0.05, **p<0.01, ***p<0.001, ****p<0.0001 (one-way ANOVA with Tukey's correction). Bars represent mean ± SD. Scale bar: 100 µm.

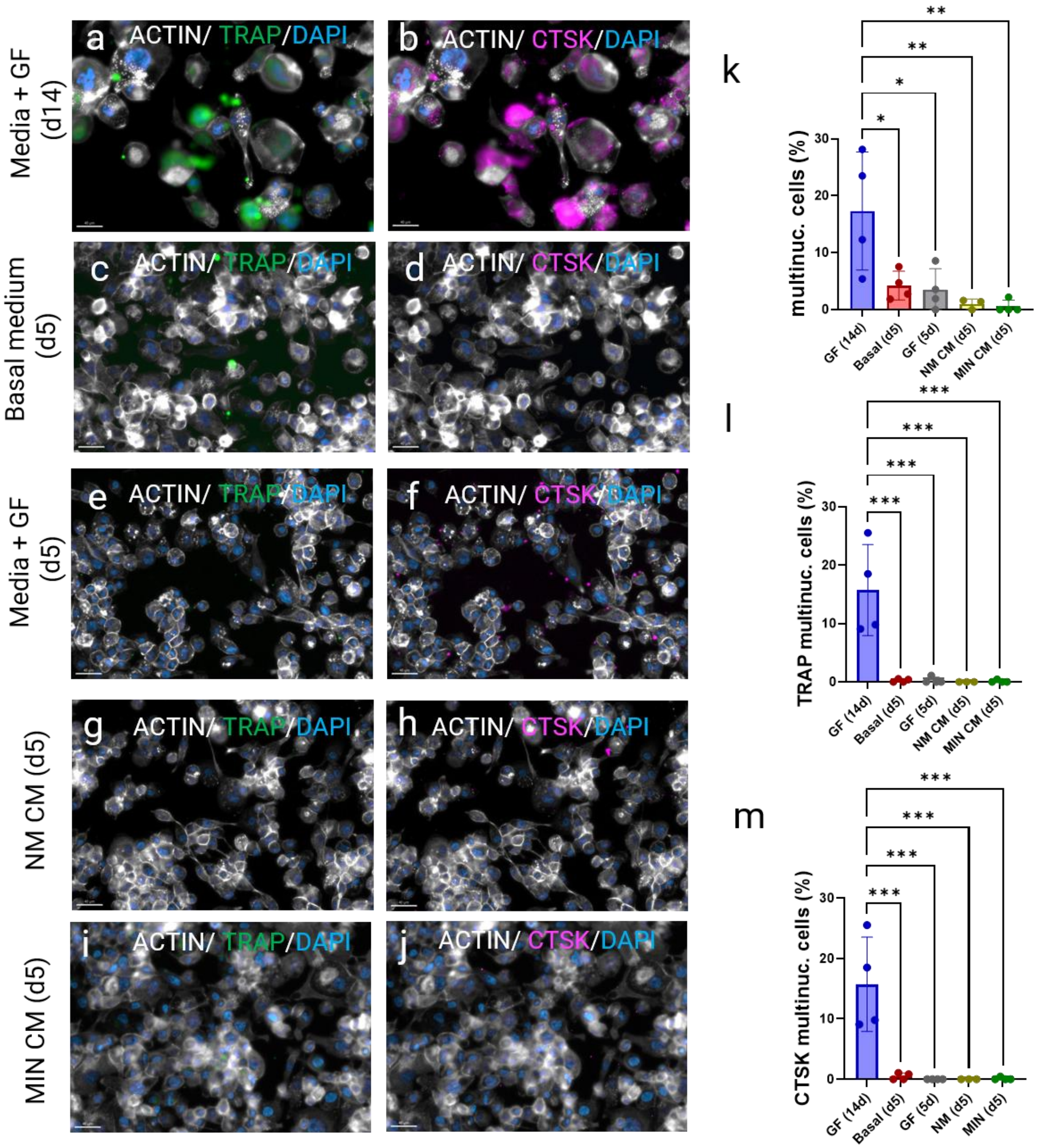


**Figure S10. Soluble factors alone are not sufficient to drive osteoclastogenesis.** Conditioned media were collected from non-mineralized (g, h) or mineralized (i, j) chips and applied to THP-1-derived macrophages (differentiated with PMA at 100 ng/mL for 48 h) for 5 days, matching the osteoclastogenesis window used in the on-chip experiments. Controls consisted of macrophages left untreated (c, d) or treated with recombinant RANKL (50 ng/mL) and M-CSF (30 ng/mL) for 5 (e-f) or 14 days (a, b). Immunofluorescence for TRAP (green) and cathepsin-K (purple), with nuclei counterstained by DAPI (blue), shows multinucleated osteoclasts predominantly following 14 days of RANKL/M-CSF treatment, with minimal multinucleation observed following conditioned medium exposure. Scale bars, 40 µm (a–j). Bar graphs (k–m) show the percentage of multinucleated cells (≥3 nuclei), the percentage of TRAP-positive multinucleated cells, and the percentage of cathepsin-K-positive multinucleated cells, respectively. Data are from 4 independent replicates; statistical significance in k–m is indicated by *p < 0.05, **p < 0.01, and ***p < 0.001, one-way ANOVA with Tukey's post hoc correction. Bars represent mean ± SD.

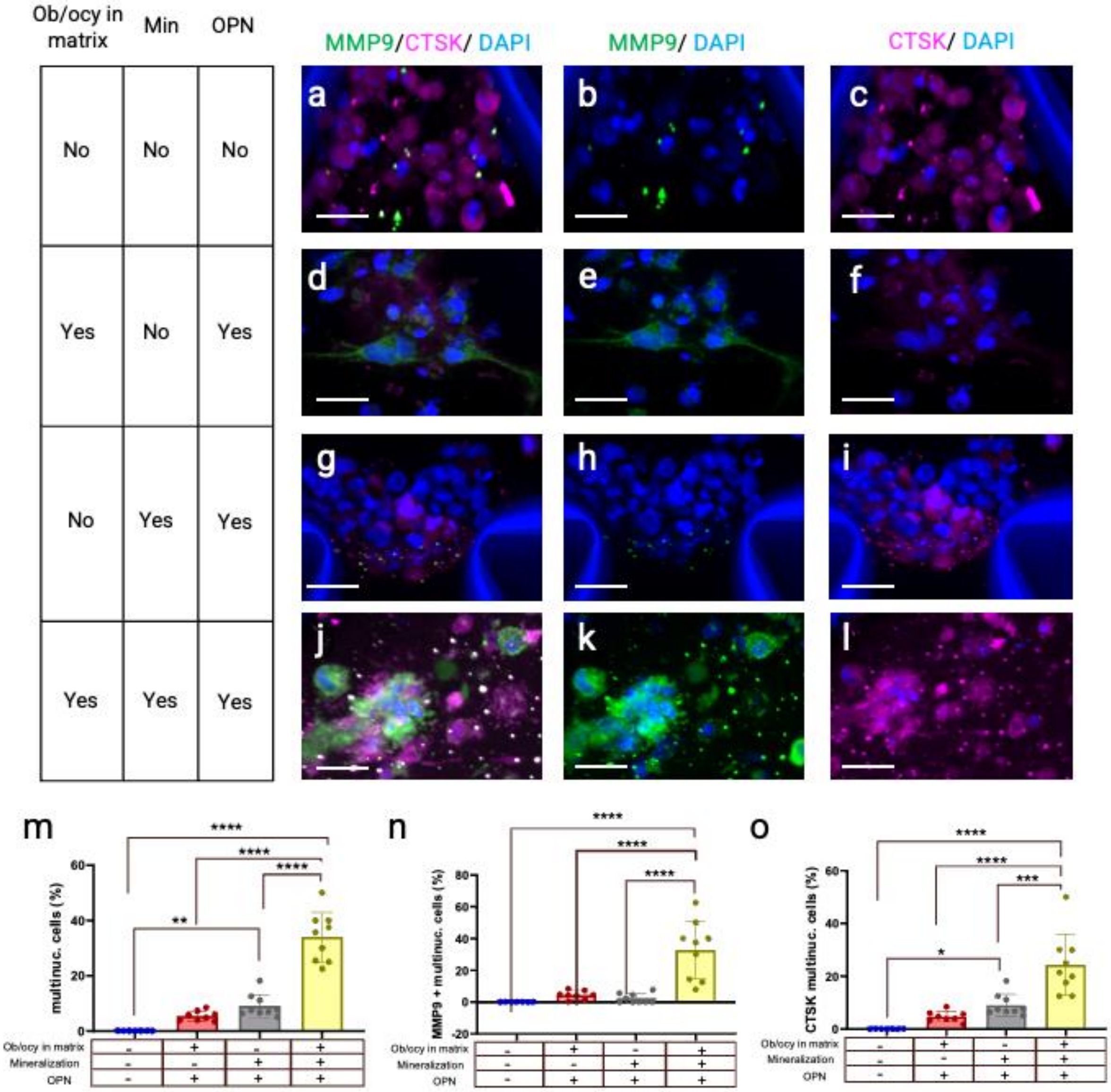

**Figure S11. The intact mineralized cellular niche promotes osteoclastogenesis beyond the effects of osteopontin and mineralized matrix.** To separate the contribution of osteopontin (OPN) from that of the mineralized matrix and its embedded cells, four conditions were compared: non-mineralized collagen without embedded cells or OPN (a–c), non-mineralized collagen containing embedded cells and supplemented with OPN (d–f), acellular collagen mineralized with OPN (g–i), and the complete construct of mineralized collagen with embedded cells and OPN (j–l). THP-1-derived macrophages were seeded into the lateral channel in all conditions and cultured for 5 days, then stained for MMP9 (green) and cathepsin-K (purple), with nuclei counterstained by DAPI (blue). Scale bars, 40 µm (a–l). Bar graphs show the percentage of multinucleated cells with three or more nuclei (m), the percentage of MMP9-positive multinucleated cells (n), and the percentage of cathepsin-K-positive multinucleated cells (o) over all cells in the lateral channel and interface from three biological replicates. Although MMP9 and CTSK expression was observed in some control conditions, marker positivity alone was not considered evidence of osteoclastogenesis, which was quantified for double positivity of marker expression and associated multinucleated cells with more than 3 nuclei. Osteoclastogenesis was observed most prominently in the complete mineralized construct (j–l), whereas the OPN-supplemented non-mineralized and acellular mineralized conditions exhibited substantially lower osteoclast-associated differentiation. These results indicate that neither condition alone was sufficient to reproduce the response observed in the complete construct. Statistical significance is indicated by $^{**}p < 0.01$, $^{***}p < 0.001$, and $^{****}p < 0.0001$, one-way ANOVA with Tukey's post hoc correction. Bars represent mean ± SD.

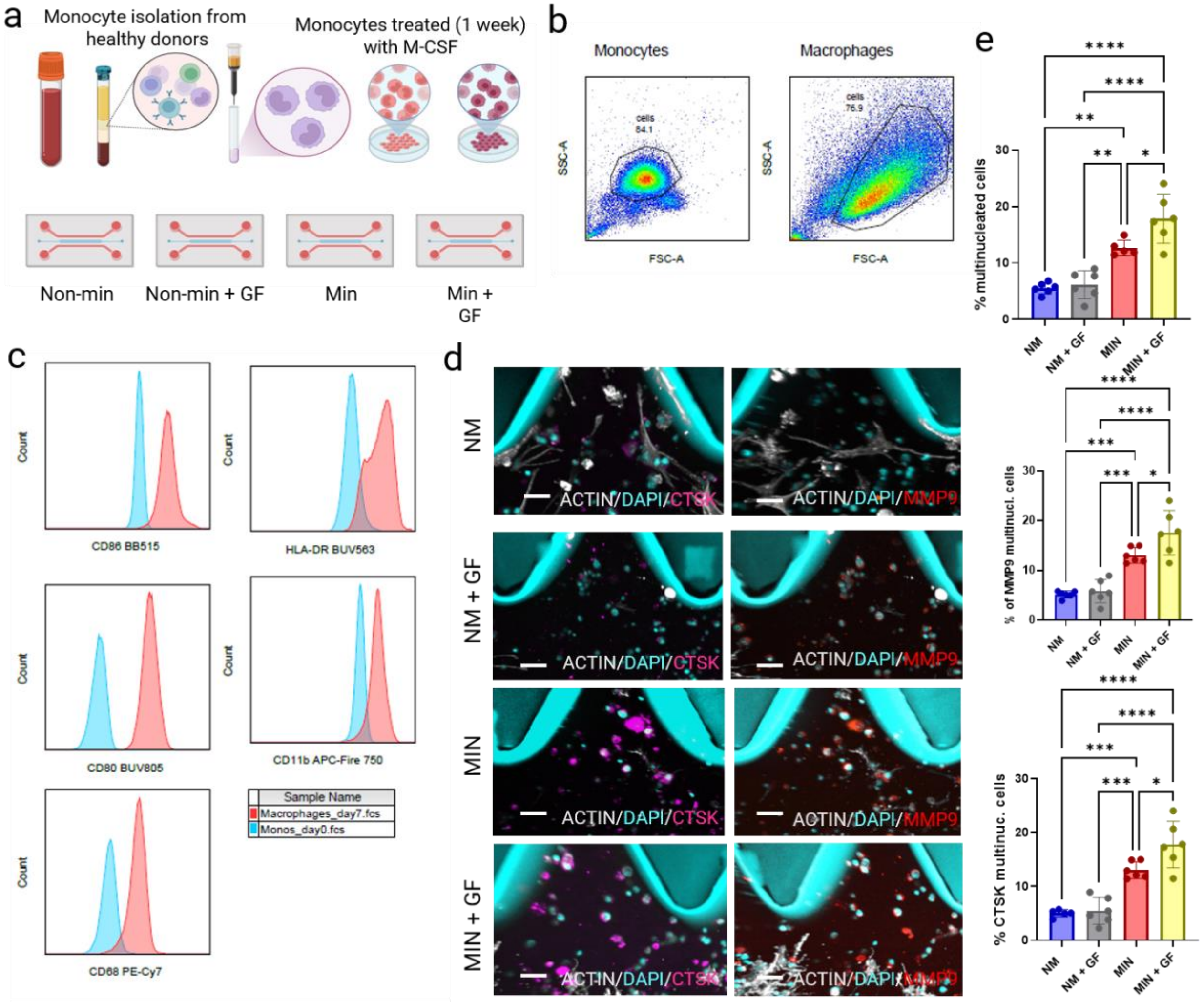


**Figure S12. Osteoclastogenesis on-a-chip using primary human cells.** Primary human monocytes were isolated from 3 different healthy donors (a) and differentiated into macrophages with M-CSF for 7 days off-chip. Flow cytometry dot plots (b) show changes in cell size and granularity following differentiation, while histograms (c) confirm distinct surface marker expression between monocytes and macrophages (CD86, HLA-DR, CD80, CD11b, CD68). Differentiated macrophages were seeded into the lateral channel of the chip and cultured for 5 days, followed by fixation and staining for cathepsin K (CTSK) and matrix metalloproteinase-9 (MMP9). Representative images (d) show increased CTSK- and MMP9-positive cells under mineralized (MIN) and mineralized plus growth factors (MIN + GF) conditions. Quantification (e) shows higher proportions of multinucleated cells and MMP9-positive and CTSK-positive multinucleated cells in MIN and MIN + GF groups. Scale bar, 50 µm. Statistical analysis was performed using one-way ANOVA with Tukey's correction (* $p < 0.05$, ** $p < 0.01$, *** $p < 0.001$, **** $p < 0.0001$). Bars represent mean ± SD.

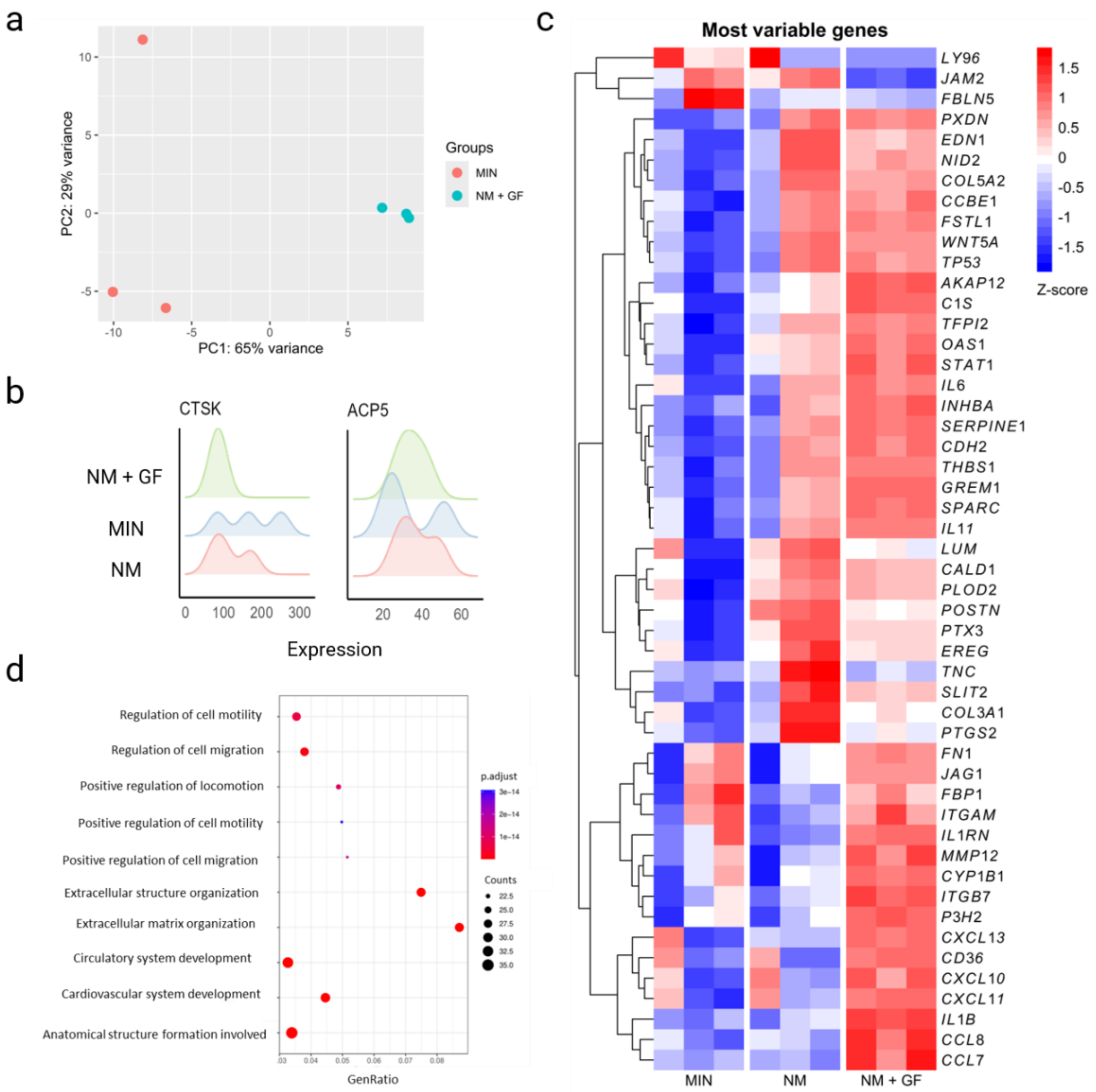


**Figure S13. Transcriptional profiling of osteoclastogenesis in mineralized and growth factor-supplemented bone-on-a-chip cultures.** (a) Principal component analysis comparing the group treated with RANKL/M-CSF (NM + GF, green dots) with the mineralized group (MIN, orange dots). Each dot represents a different replicate, and the data show a clear separation between the two groups, with the replicates of the NM + GF group clustering closely together. (b) Ridge plot showing the expression of the known osteoclast marker genes CTSK and ACP5. (c) Heatmap of the most variable genes for the mineralized (MIN), non-mineralized (NM), and non-mineralized with RANKL/M-CSF (NM + GF) groups. Color intensity represents the Z-score, with red indicating higher expression and blue indicating lower expression across three biological replicates. (d) Gene ontology (GO) enrichment analysis of differentially expressed genes in the mineralized group (MIN) compared with the RANKL/M-CSF group (NM + GF).

The y-axis represents gene functions, the x-axis represents the gene ratio, and dot size indicates the number of genes associated with each enriched term; dot color represents the adjusted p-value.

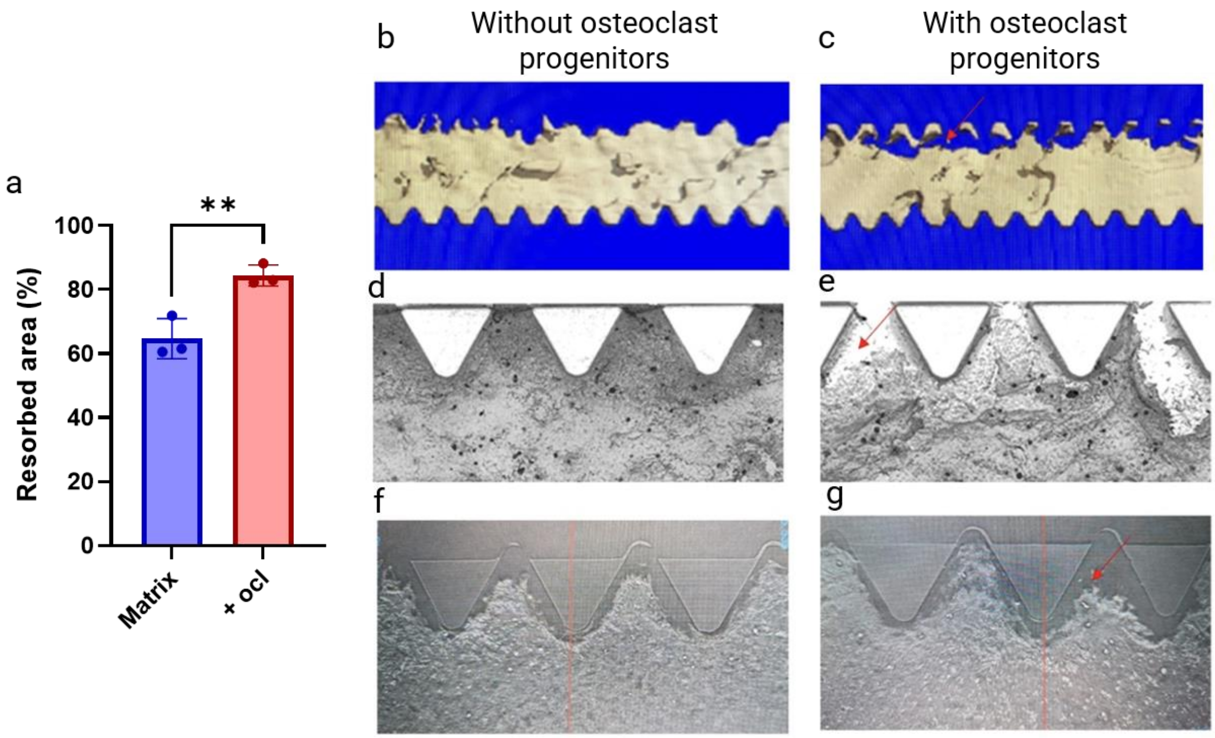


**Figure S14**. **Bone-on-a-chip–generated osteoclasts exhibit functional matrix resorption**. (a) Quantification of resorbed matrix area in mineralized constructs with (+Ocl) or without (-Ocl) osteoclast progenitors. Representative 3D micro–CT reconstructions and corresponding images of the mineralized matrix and channel interface of mineralized chips cultured without (b, d, f) or with osteoclast progenitors (c, e, g) reveal localized matrix loss and interfacial cavities in progenitor-containing conditions (red arrows). Bars represent mean ± SD.

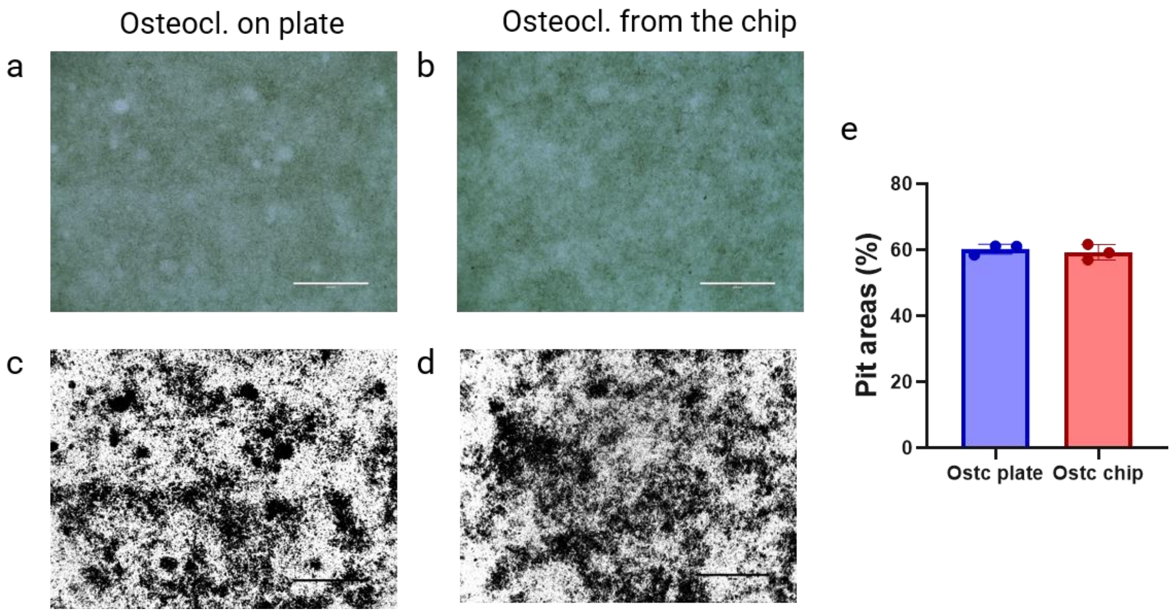


**Figure S15. Chip-derived osteoclasts are functional on CaP plates.** Functional activity of osteoclasts differentiated on mineralized matrices in chips before being retrieved and seeded on CaP plates for 2 days (b,d) was compared to that of osteoclasts derived from macrophages cultured on CaP plates for 14 days in media supplemented with RANKL (50 ng/mL) and M-CSF (30 ng/mL) for 14 days (a,c). Resorbed pit area fraction (e) quantified from bright-field images converted to high-contrast binary images in ImageJ according to AMSBIO (Life Science and Product Solutions) guidelines showed no significant differences between osteoclasts differentiated directly on the plate and osteoclasts from the chip. Bars represent mean ± SD.

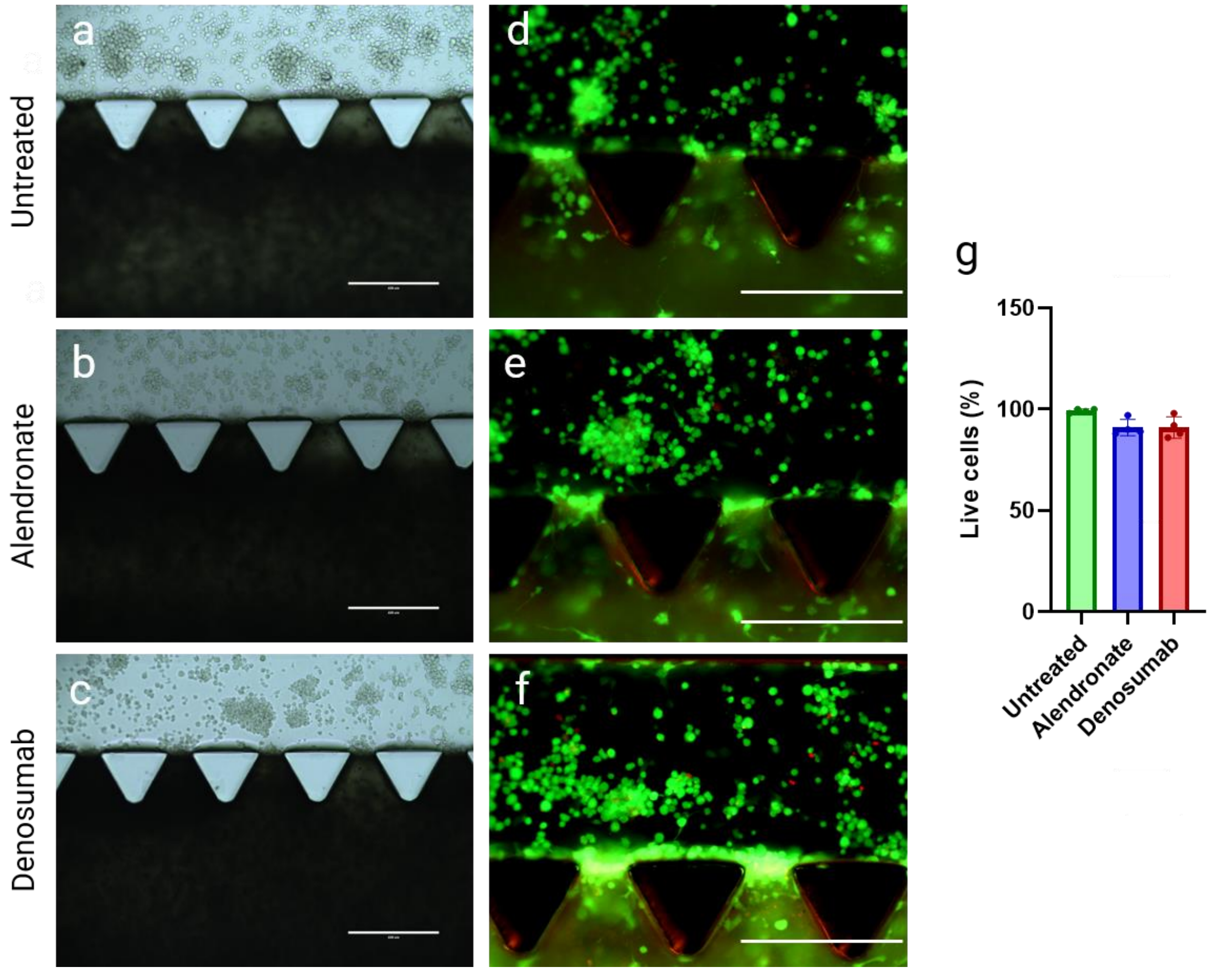


**Figure S16. Cellular viability after drug treatment**. Representative phase contrast images (a-c) of mineralized chips where macrophages received no treatment (a), alendronate (b), or denosumab (c) showed comparable cell numbers. Cell viability assessed by calcein AM (live, green) and propidium iodide (dead, red) staining (d-f) showed (g) no significant differences in cell viability between the groups (N = 3) after 5 days of treatment. Scale bars represent 400 µm. Bars represent mean ± SD.

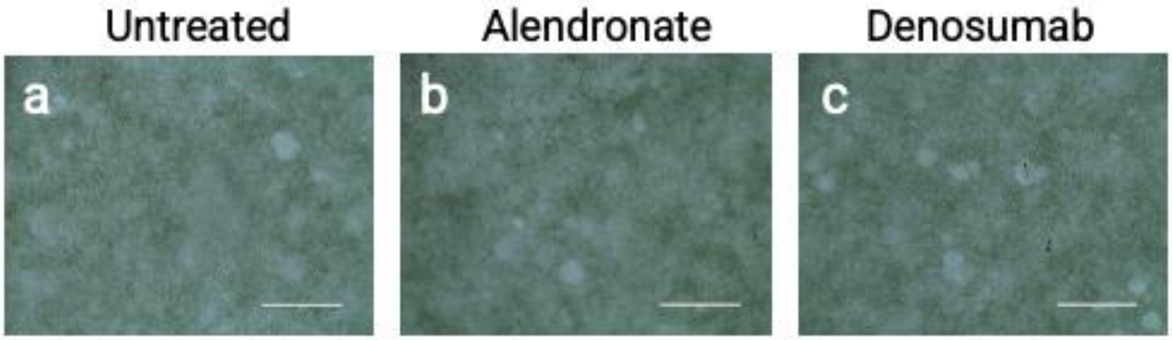


**Figure S17. Functional assessment of osteoclasts differentiated on CaP plates following drug treatment.** Representative resorption pits on CaP-coated plates following 14 days of culture of macrophages in medium supplemented with RANKL (50ng/mL) and M-CSF (30 ng/mL), either without drug treatment (a), or with alendronate (b) or denosumab (c). Panels (a-c) show phase-contrast images of the samples (N = 3). Scale bars represent 400 μm.

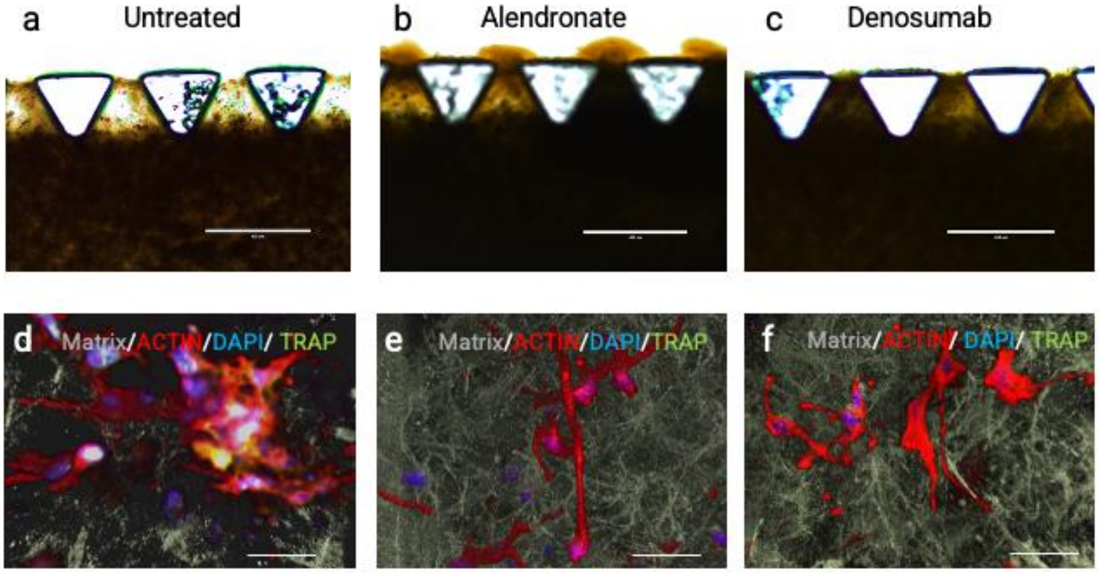


**Figure S18. Alizarin red and confocal image screening of bone resorption on bone-on-a-chip treated with anti-resorption drugs.** Alizarin red staining of bone-mimetic environments shows higher resorption of mineralized matrix in untreated samples (a) when compared to samples treated with alendronate (b) or denosumab (c), demonstrating the ability of the model to capture drug-specific effects on bone resorption in vitro. Scale bars (a-c): 400 µm. Panels (d–f) show cell–matrix interactions with actin, DAPI, and TRAP staining in untreated (d), alendronate-treated (e), and denosumab-treated (f) samples, revealing reduced TRAP staining in both drug-treated groups. Scale bars (d-f): 40 µm

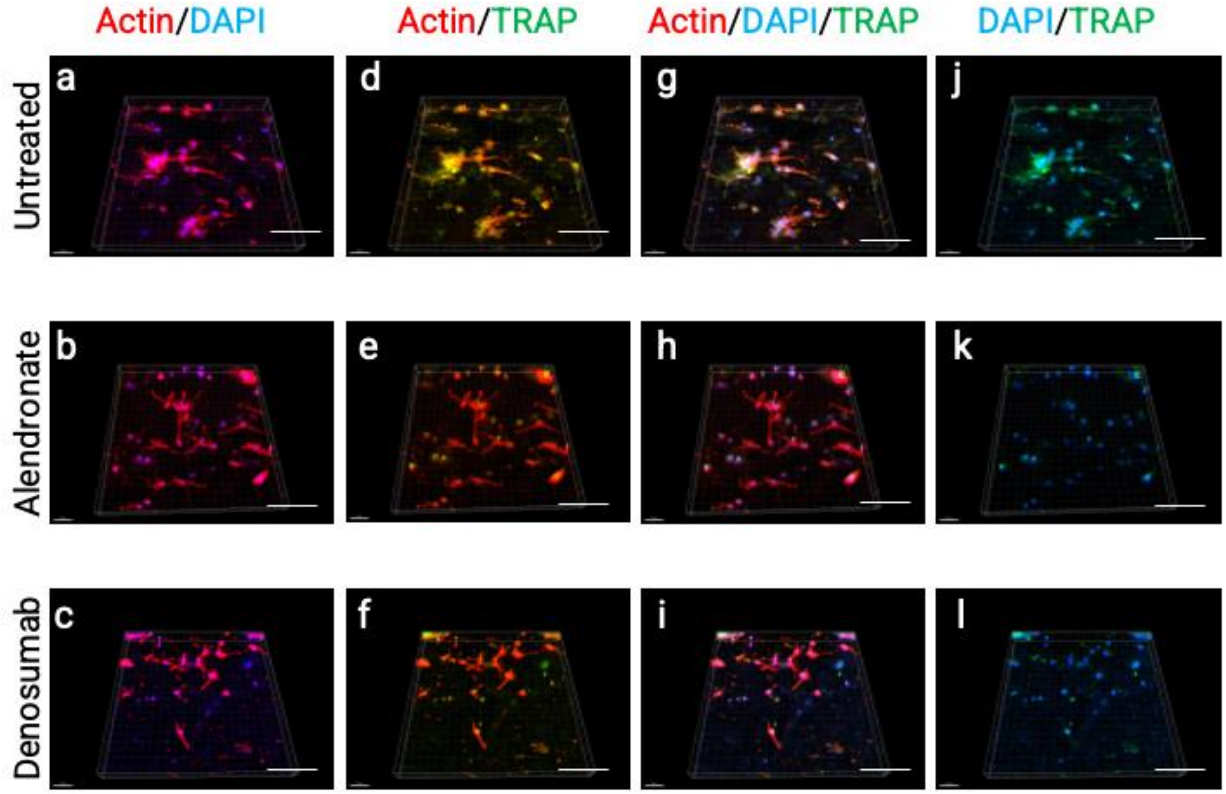


**Figure S19. Bone-on-a-chip models osteoclast-targeting drug responses through reduced TRAP staining.** Representative lower-magnification images of mineralized chips treated with alendronate (b, e, h, k) or denosumab (c, f, i, l) for 5 days compared to the untreated group (a,d, g, and j) and immunostained for actin, DAPI, and TRAP (N=3). Reduced TRAP staining in drug-treated groups, alendronate (b, e, h, k) and denosumab (c, f, i, l), highlighting the platform's capacity to model osteoclast-targeting drug responses. The scale bars represent 150 µm.

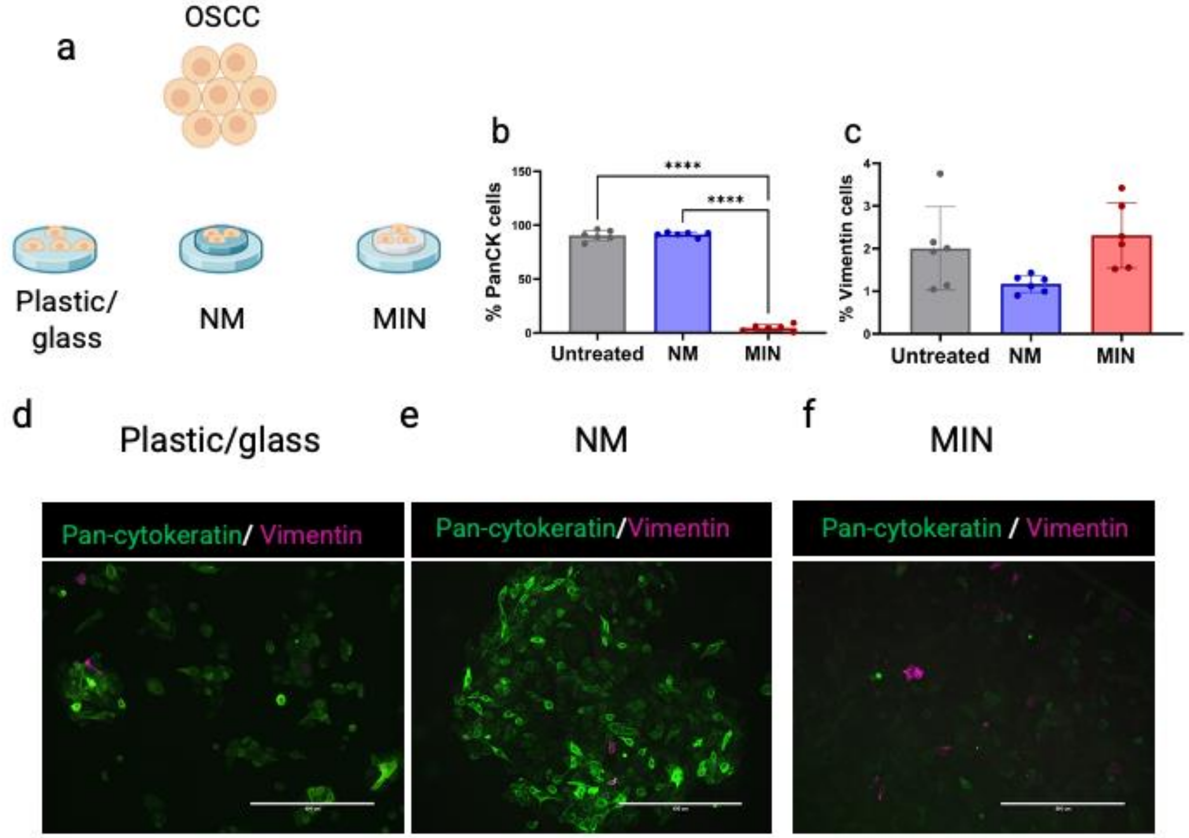


**Figure S20. Mineralized matrix promotes loss of Pan-cytokeratin in OSCC cells off chip**. UCSF-OT-1109 OSCC cells were seeded onto collagen disks that were either mineralized or non-mineralized, as well as on tissue culture plastic. After 3 days, samples were fixed and immunostained for pan-cytokeratin (PanCK, green) and vimentin (magenta) (a). Quantification (b) reveals a significant reduction in PanCK positivity in cells on mineralized collagen (f) compared with non-mineralized collagen (e) or plastic (d). Scale bars in (d–f) represent 400 µm. Statistical significance was determined by one-way ANOVA with Tukey's post hoc test (****$p < 0.0001$).

**Supplementary methods:**

**Computational fluid dynamics modeling**

The bone-on-chip geometry was designed in Autodesk Fusion 360 (Autodesk, San Rafael, CA) and exported as a STEP file, which was imported into COMSOL Multiphysics (v6.3, COMSOL AB, Stockholm, Sweden) to build a three-dimensional model of oscillatory perfusion in the media channels and interstitial flow through the central collagen matrix. The Free and Porous Media Flow (Brinkman) interface was used, with the media channels defined as free-flow domains and the central collagen compartment defined as a homogeneous, isotropic porous domain (porosity $\varepsilon = 0.99$, permeability $k = 1.0 \times 10^{-12}$ m². The culture medium was assumed to be an incompressible Newtonian fluid with a density of 1000 $kg \cdot m^{-3}$ and a dynamic viscosity of 0.001 Pa·s, and a no-slip condition was imposed on all channel and chamber walls.

The flow generated by platform rocking was represented as a time-dependent hydrostatic pressure difference between the surfaces of the two reservoirs, given by $\Delta P(t) = \rho g L \cdot \sin(\theta_0 \sin(2\pi f t))$. In this expression, ρ is the fluid density, g is the gravitational acceleration, L is the center-to-center distance between the reservoirs, $\theta_0$ is the maximum rocking angle (15°), and f is the rocking frequency (1 Hz). To drive reciprocating flow, pressures of $+\Delta P(t)/2$ and $-\Delta P(t)/2$ were prescribed on the two reservoir surfaces.

The oscillatory flow field was resolved with a transient study spanning multiple rocking cycles, which provided the pressure distribution, velocity magnitude, and flow direction throughout the device. Mean velocities in the media channels and in the collagen matrix were calculated by averaging over the corresponding domains. The volumetric flow rate through the media channels was computed as $Q = \int_A u \cdot n \, dA$, where A is the channel cross-sectional area, u is the velocity vector, and n is the outward unit normal. The hydrodynamic stimulus experienced by cells within the collagen matrix was estimated as an effective cell-scale shear stress, $\tau = \mu U/\sqrt{k}$, where μ is the dynamic viscosity, U is the local interstitial velocity obtained from the Brinkman solution, and k is the collagen permeability (Supplementary Fig. S6).

**Cellular viability**

After 3 days of mineralization followed by 5 days of osteoclastogenic co-culture (day 8), the cells inside the matrix were stained with a Live/Dead Cell Imaging Kit (Molecular Probes, Thermo Fisher) to confirm their viability. Following incubation, the specimens were washed with PBS and imaged using an inverted fluorescence microscope (n = 4) (FL Autos, EVOS – Thermo Fisher Scientific, Waltham, MA, USA). Image processing and analysis were performed using ImageJ (FIJI) software87 to assess cell viability, calculated as the proportion of live cells (total DAPI-stained nuclei minus propidium iodide (PI)-positive dead cells) relative to the total number of DAPI-stained nuclei, expressed as a percentage.

**Conditioned-medium osteoclastogenesis assay**

To test whether soluble factors secreted by the bone-on-a-chip constructs were sufficient to drive osteoclastogenesis independently of the on-chip microenvironment, conditioned medium was collected from the main channel of non-mineralized and mineralized chips after 3 days of culture. Collected medium was filtered through a 0.22 µm filter to remove debris and used fresh.

THP-1 monocytes were seeded in 96-well plates at ($3 \times 10^4$ cells/well) and differentiated into macrophages with PMA (100 ng/mL) for 48 h. Macrophages were then cultured for 5 days under four conditions: (i) macrophages alone (negative control), (ii) macrophages treated with RANKL (50 ng/mL, PeproTech) and M-CSF (30 ng/mL, PeproTech), (iii) conditioned medium from non-mineralized chips, and (iv) conditioned medium from mineralized chips. Conditioned medium was applied and replenished every 3 days. In a separate experiment, macrophages were maintained with RANKL (50 ng/mL) and M-CSF (30 ng/mL) for 14 days to confirm osteoclastogenic competence under prolonged stimulation (positive control).

After culture, cells were fixed in 10% neutral-buffered formalin for 30 min at room temperature and stained for TRAP (Novus Biologicals, NBP2-45294, 1:100) and cathepsin K (Thermo Fisher, PA5-18950, 1:100) as described. Plates were imaged using a Zeiss Apotome 3 on an Axio Observer. Osteoclast differentiation was quantified as the number of TRAP-positive multinucleated cells (≥3 nuclei) per well (3 field/view) (N = 4).

**Osteopontin control experiment**

To determine whether osteopontin or the mineralized matrix alone was sufficient to induce osteoclastogenesis, independently of cells embedded within the matrix, four conditions were compared: (i) non-mineralized collagen without OPN, (ii) non-mineralized collagen supplemented with OPN (100 µg/mL Lacprodan OPN-10), (iii) acellular mineralized collagen (mineralized in the absence of embedded osteoblasts), and (iv) mineralized collagen containing embedded osteoblasts (complete construct). Mineralized conditions were prepared using the standard mineralizing medium (9 mM $CaCl_2{\cdot}2H_2O$, 4.2 mM $K_2HPO_4$, 100 µg/mL Lacprodan OPN-10, 25 mM HEPES) replaced daily for 3 days, as described above. Osteoblast-laden constructs were prepared at $3 \times 10^6$ cells/mL as described; acellular constructs were prepared identically but without cells.

THP-1-derived macrophages were then seeded into the lateral channel ($3 \times 10^4$ cells) and cultured for 5 days. Samples were fixed in 10% neutral-buffered formalin for 30 min at room temperature and stained for MMP9 (Bioss, bs-4593R, 1:100) and cathepsin K (Thermo Fisher, PA5-18950, 1:100) as described. Osteoclast differentiation was quantified as the number of multinucleated cells (≥3 nuclei), MMP9-positive multinucleated cells, and cathepsin K-positive multinucleated cells (N = 3).

**Supplementary Movie 1**. **Mineralized bone-on-a-chip supports vascular and osteoclast compartments.** Representative 3D visualization showing a mineralized bone-on-a-chip incorporating vascular and osteoclast components. Vascular networks were formed by co-culturing mesenchymal stem cells and human umbilical vein endothelial cells (HUVECs), together with osteoblasts. After 3 days of mineralization, the system exhibits lumenized vessel structures and osteocyte differentiation. Macrophages were introduced into the lateral channel and differentiated into osteoclasts. Vessels are labeled by CD31 (orange), osteocytes by PDPN (yellow), osteoclasts by cathepsin K (CTSK; purple), and nuclei by DAPI (blue).

**Supplementary Movie 2. Remodeling niches in osteocyte–osteoclast co-cultures.** This video demonstrates the dynamic remodeling regions formed within mineralized bone-like matrices following 5 days of macrophage seeding in the lateral channel. Osteoclasts (TRAP-positive, green; actin-positive, red; DAPI-positive, blue) are observed interacting directly with osteocytes (actin-positive, red; DAPI-positive, blue) embedded within the mineralized matrix (grey), imaged by second harmonic generation microscopy. These localized remodeling zones highlight the power of this platform to recapitulate osteocyte–matrix–osteoclast interactions and provide unique insights into bone remodeling processes.

**Supplementary Table S1. Gene panel used for targeted NanoString transcriptional profiling.** Complete list of genes included in the NanoString PanCancer Progression Panel supplemented with 55 custom bone-associated genes, comprising 825 genes in total.
